# Cribriform Plate Microenvironment Assembles a Suppressive Myeloid Network during EAE-induced Neuroinflammation

**DOI:** 10.64898/2026.01.07.698165

**Authors:** Collin Laaker, Martin Hsu, Andy Madrid, Jenna Port, Sophia M. Vrba, Melinda Herbath, Cameron Baenen, Mohan Kumar, Thanthrige Thiunuwan Priyathilaka, Matyas Sandor, Zsuzsanna Fabry

## Abstract

During neuroinflammation, CD11c+CD11b+ myeloid cells accumulate at the cribriform plate, a key cerebrospinal fluid (CSF) and antigen outflow site in mice. At this site, podoplanin (PDPN)-expressing cells, including lymphatic vessels and meningeal layers, expand to create a distinct drainage microenvironment. In this study we sought to characterize myeloid cells which populate this region using a mouse model of neuroinflammation, experimental autoimmune encephalomyelitis (EAE). Utilizing a combination of immunohistochemistry, flow cytometry, and scRNAseq, we report that macrophages and dendritic cells (DCs) from this region display unique expressional signatures related to tolerance, cell death, and reduced inflammatory profile. Together this data supports that myeloid retention at the cribriform plate and olfactory bulb meninges promotes a local immunosuppressive environment.

## Introduction

Meningeal microenvironments are highly dynamic during neuroinflammatory disease, which is critical for the progression and resolution of inflammation in the central nervous system (CNS) (Rustenhoven et al., 2021; Hitpass Romero et al., 2025). For example, in the dura, the outermost meningeal layer, lymphatic vessels can adapt and respond to brain inflammation; sampling cerebrospinal fluid (CSF), waste, and CNS-antigens to help coordinate immunity during neuroinflammation (Zhang et al., 2025b; Louveau et al., 2018). Additionally, CD11b^+^CD11c^+^ peripheral myeloid populations rapidly infiltrate the brain and meningeal niches, accessing antigen, upregulating MHC-II, and migrating through this meningeal lymphatic network, underscoring their role in antigen presentation and T-cell activation during infection and autoimmunity in the central nervous system (CNS) (Jordão et al., 2019; Clarkson et al., 2015, 2017; Louveau et al., 2018). While there are increasing studies on the regulation of immune cells in the dorsal dural immune hubs (Rustenhoven et al., 2021; Fitzpatrick et al., 2024), there is still little knowledge of the phenotypes of myeloid cells that accumulate at CSF efflux sites along the base of the brain, like the cribriform plate lymphatics.

Recently, our lab and others have shown that Lyve-1^+^ lymphatic vessels between the olfactory bulbs access CSF through gaps in the arachnoid layer along olfactory nerve bundles, enabling fluid, antigen, and immune cell drainage to the cervical lymph nodes (Hsu et al., 2019, 2022; Spera et al., 2023; Jin et al., 2025). This lymphatic niche at the cribriform plate expands during experimental autoimmune encephalomyelitis (EAE), a mouse model of Multiple Sclerosis (MS), and upregulates genes related to leukocyte cross-talk (adhesion, chemotaxis, antigen processing and presentation, and activation) (Hsu et al., 2022). Furthermore, we previously showed that a majority of the retained cells at the cribriform plate lymphatic niche during neuroinflammation were CD11c^+^ CD11b^+^ (Hsu et al., 2019, 2022), implicating myeloid subsets, including DCs, were the primary immune cells that are recruited, retained, and potentially regulated in the meningeal-lymphatic environment at the cribriform plate(Hsu et al., 2022).

In the context of brain inflammation, the regulation and efflux pathways of DCs from the CNS are still poorly understood (Laaker et al., 2023). Our lab and others have shown that during EAE, intracerebrally and intracisternally injected DCs can exit the intracranial space and drain to the cervical lymph nodes in a CCR7-dependent manner (Clarkson et al., 2017; Karman et al., 2004; Louveau et al., 2018). Interestingly, intracranially retained CCR7-KO DCs exacerbate autoimmune-mediated neuroinflammation, restimulating infiltrating autoreactive T-cells near brain tissue (Clarkson et al., 2017). While evidence supports that effluxing pro-inflammatory DCs from the intracranial space can further stimulate T-cell populations in the draining lymph nodes (Hsu et al., 2019), there is also data to support that DCs migrating in the olfactory bulb region of mice acquire a tolerogenic cell state (Mohammad et al., 2014). Thus, understanding how CD11b^+^CD11c^+^ cells interact at the cribriform plate could give important insight into how DC cell states are conditioned at CSF-efflux from the skull. Importantly, cranial nerve bundles, like the brain, are also surrounded by meningeal layers and interface with CSF to form a specialized perineural microenvironment (PME) as they extend from the CNS and traverse the skull (Fahmy et al., 2021; Proulx, 2021).

While peripheral nerve bundles are increasingly recognized as a unique neuroimmune niche that promote immunosuppressive tumor microenvironments (TME)(Baruch et al., 2025; Liebig et al., 2009; Chen et al., 2019) and tissue repair after injury through reprogramming of local and recruited myeloid cells(Ydens et al., 2020), the PME of cranial nerve bundles as they exit the skull during neuroinflammation is still poorly understood. Further complicating analysis, inflammatory environments in the meninges, including near the PME, often elevate extracellular matrix (ECM)-related tissue niches where immune cells, lymphatics, fibroblasts, and blood vessels are tightly interwoven in dense cell-cell interactions (Dorrier et al., 2022). Interestingly, during EAE we find increased podoplanin expression at the cribriform plate which is rich with myeloid cells. Understanding this cell-cell cross-talk could be essential to understanding how DCs and other myeloid cells contacting the meningeal-lymphatic environment are reprogrammed during disease.

In this study, we identified populations of CCR7^+^ DCs within this microenvironment that had enriched pathways related to programmed cell death (PD-1), tolerance induction, and inhibited interferon-signaling pathways. Additionally, our scRNAseq dataset and IHC analysis support for a role of CCR2^+^ monocytes and integration of alternatively activated Arginase-1 (*Arg1*) and CHI3L3 (*Chil3*) macrophages into a PDPN^+^ microenvironment at the cribriform plate during EAE. We hypothesize that the cribriform plate functions as an immune interface between the CNS and the periphery, simultaneously accessing draining antigen and CSF while dynamically reshaping its perineural myeloid microenvironment to regulate the phenotype of DCs and antigen outflow to the cervical lymph nodes.

## Results

### CSF1R^+^, CD11c^+^, and MHC-II^+^ cells are in cell-cell contact with an expanded PDPN^+^ PME around olfactory nerve bundles during EAE

We previously identified that CD11b^+^ CD11c^+^ myeloid cells increase near olfactory nerves and along cribriform lymphatic vessels during EAE-induced inflammation (Hsu et al., 2022; Laaker et al., 2025). To more holistically understand the spatial environment of myeloid cells at the meningeal–lymphatic drainage regions of the cribriform plate (**Figure 1A**), we induced EAE in CSF1R-GFP mice and stained decalcified cribriform plate sections for the common lymphatic markers PDPN and LYVE-1 (**Figure 1B**). As reported previously (Hsu et al., 2019), LYVE-1^+^ regions are primarily found in the spaces directly between the olfactory bulbs, running adjacent to outwardly projecting midline olfactory nerves above the cribriform plate (**Figure 1B**). Directly surrounding olfactory nerve bundles is a prominent PDPN^+^LYVE-1^neg^ layer which links the perineural region to double positive PDPN^+^LYVE-1^+^ lymphatics (**Figure 1B; zoom panel**).

**Figure 1.**
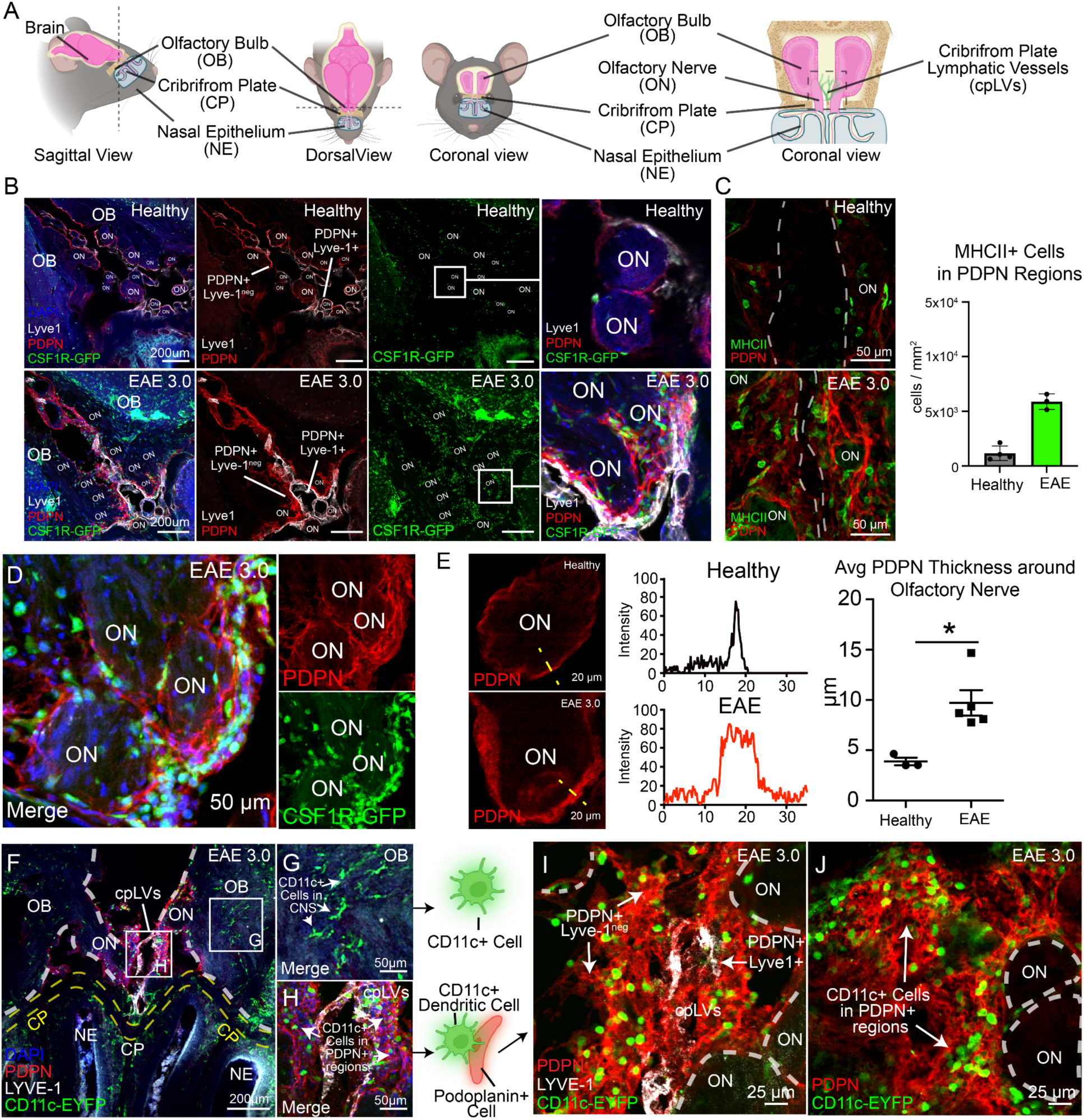
Podoplanin expands and interacts with myeloid cells at the cribriform plate during EAE. **(A):** Cartoon schematic of the cribriform plate lymphatic vessels (cpLVs) which associate the perineural environment of olfactory nerves (ON) between the olfactory bulbs (OB) in mice. Lymphatic vessels are directly next to ONs as they project through the cribriform plate (CP) into the nasal epithelium (NE). **(B):** Healthy and EAE 3.0 CSFR1R-GFP mouse CP sections were stained for podoplanin (PDPN) PE and Lyve-1 Alexa 660 to visualize the relationship of myeloid cells (CSF1R-GFP+), lymphatics (Lyve-1+, PDPN+), and meningeal stroma (Lyve-1_negative_, PDPN+). Zoomed panels show the perineural immune-stromal-lymphatic environment around ONs during healthy and EAE. **(C):** Quantification of MHCII MFI at the CP during healthy and EAE 3.0 conditions using identical imaging conditions. MHCII (eFlour 450) increases in PDPN+ regions during EAE. Unpaired student’s t-test. **P < 0.01. Data are represented as mean ± SEM. **(D):** Representative image of perineural CSF1R-GFP cells adhered to PDPN+ region of ONs. **(E):** Quantification of average PDPN+ thickness (µm) around ONs during healthy and EAE conditions. Intensity plot profiles (yellow dotted line) were used to estimate PDPN+ regions. n=3-5 mice. *P<0.05. Data are represented as mean ± SEM **(F):** Representative image of immune response at cribriform plate in EAE 3.0 CD11c-eYFP mice stained for Lyve-1 and PDPN. **(G):** Non-Lymphatic (PDPN_negative_, Lyve1_negative_) associated CD11c+ dendritic cells are in the olfactory bulb brain tissue. The cells likely represent infiltrating DCs, macrophages, and activated Microglia. **(H):** Meningeal**-**Lymphatic (PDPN^+^) associated CD11c+ are at the cribriform plate lymphatics. The cells likely represent infiltrating DCs, macrophages, and monocytes **(I-J):** Representative image of immune response at cribriform plate in EAE 3.0 CD11c-eYFP mice stained for Lyve-1 and PDPN. CD11c-eYFP+ embed into dense PDPN+ ECM) near cribriform lymphatics (Lyve-1).

PDPN not only labels lymphatic vessels, but also efficiently labels meningeal layers surrounding them along the olfactory nerve layer, including fibroblasts and their associated extracellular matrix (ECM) (Hitpass Romero et al., 2025; Møllgård et al., 2023; Çavdar et al., 2024). To understand if this PDPN increases around the olfactory nerve layer and is associated with antigen presenting cells (APCs) during EAE, we costained for MHCII and PDPN. We found an elevated number of MHC-II^+^ cells in PDPN regions, likely DCs, activated macrophages, and B cells (**Figure 1C**). CSF1R-GFP^+^ myeloid cells were colocalized to PDPN^+^ PME during EAE (**Figure 1D**), with perineural PDPN increasing in thickness during active EAE (**Figure 1E**). To better visualize CD11c^+^ DCs, we analyzed sections from CD11c-eYFP mice during peak EAE disease severity, and observed similar relationships between DCs and perineural PDPN^+^ regions (**Figure 1F-H**). While many CD11c^+^ cells within the parenchyma of the OB are likely activated microglia (**Figure 1G**), large numbers of CD11c^+^ cells were also observed in direct contact with perineural PDPN^+^ at the cribriform plate (**Figure 1H),** and embedded into the ECM (**Figure 1I-J**). CD11c⁺ cells in the olfactory bulb parenchyma display a ramified, microglia-like morphology consistent with tissue-resident or parenchymal surveillance cells, whereas those infiltrating the cribriform plate perineural niche show a rounded, non-ramified morphology more consistent with recently recruited monocyte-derived DCs or macrophages. Additionally, while PDPN labels the cribriform plate lymphatic vasculature, it also defines the meningeal-immune interface at the border of both the olfactory bulb and olfactory nerve bundles. Thus, the PDPN^+^ PME harbors elevated numbers of myeloid cells that directly associate with the cribriform plate lymphatic vasculature and meningeal stroma.

### Cribriform plate PME is embedded with ARG1 and CHI3L3 cells during neuroinflammation

Podoplanin ECM environments are commonly associated with immunosuppressive macrophages (Wang et al., 2022a; Swartz and Lund, 2012; Bieniasz-Krzywiec et al., 2019). To determine whether these alternatively activated macrophages accumulate *in situ* at the cribriform plate during EAE, we performed immunofluorescence staining on decalcified cribriform plate sections using the alternative activation markers CHI3L3 (*Chil3*) and ARG1 (*Arg1*), together with Lyve-1 and PDPN to delineate cribriform meningeal lymphatic structures. Here, we found CHI3L3^+^ cells increased at the cribriform plate during EAE (**Figure 2A-E**), with many CHI3L3^+^ cells adhering directly to the PDPN^+^ PME regions (**Figure 2B-C**). Similarly, Arg1-expressing cells at the cribriform plate increased during EAE and were highly colocalized within perineural PDPN^+^ (**Figure 2F-H**). However, as the size of PDPN regions increases, subsequently the regions which can hold ARG1+ and CHI3L3+ cells also increase making interpretation difficult. Together, our data suggests the accumulation of alternatively polarized macrophages in PDPN^+^ PME regions during peak EAE disease severity (Murray, 2017; Jiang et al., 2014; Xin et al., 2025).

**Figure 2.**
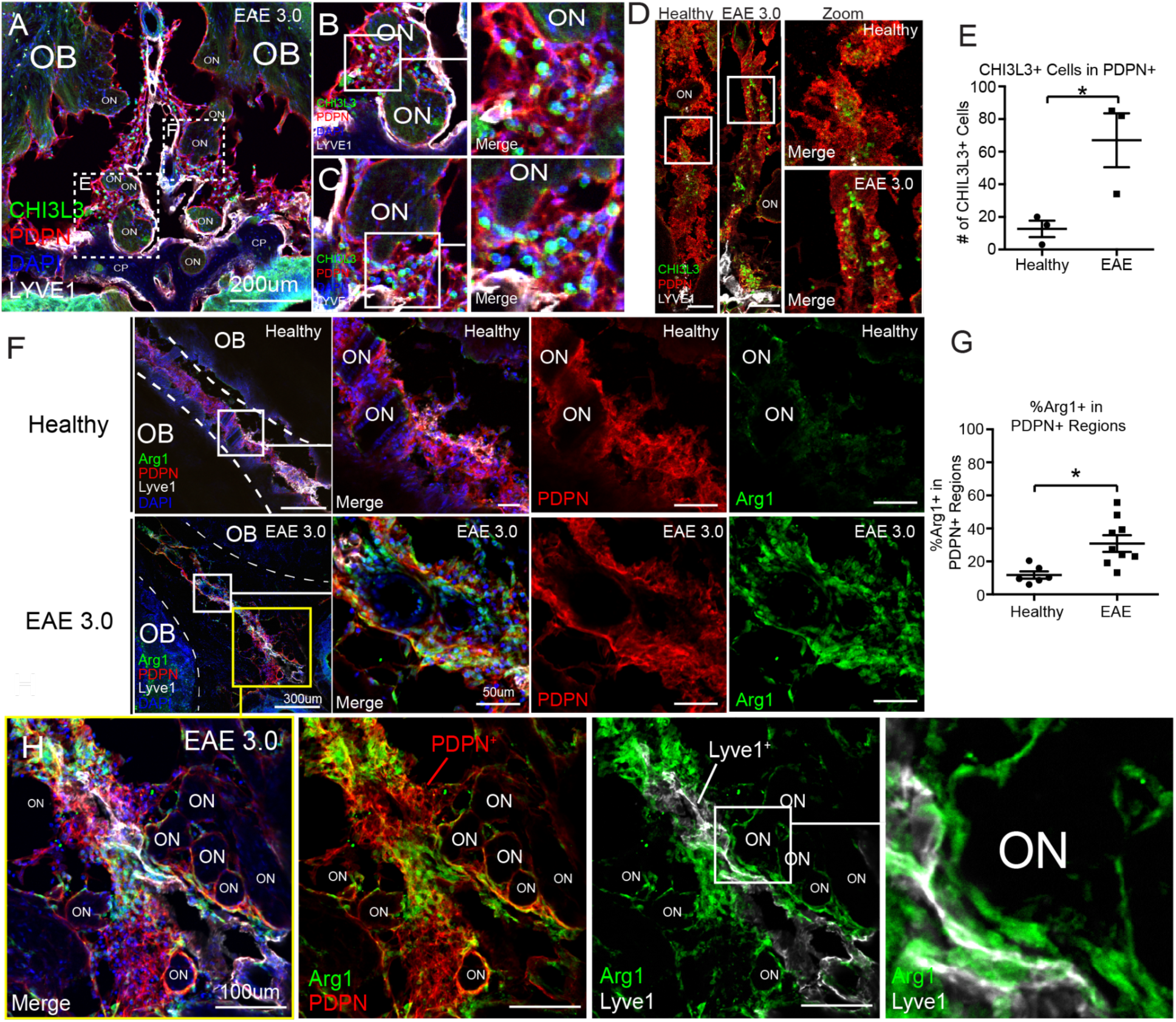
Elevation of CHI3L3 and ARG1 at cribriform lymphatics during EAE. **(A-C):** Magnification between an EAE 3.0 mouse olfactory bulb (OB) at cribriform plate (CP) showing infiltration of CHI3L3⁺ cells (green) surrounding olfactory nerves (ON) and localizing within PDPN⁺ stroma networks (red) and LYVE-1⁺ lymphatic vessels (white). DAPI marks nuclei (blue). (**B-C**) High-magnification views of boxed regions showing accumulation of CHI3L3⁺ cells in proximity to ONs and embedded within PDPN⁺ stromal-lymphatic regions and LYVE-1⁺ lymphatic zones. **(D-E)**: (**D**) Representative confocal images of healthy and EAE 3.0 tissues showing increase of CHI3L3⁺ cells (green) in PDPN⁺ regions during inflammation. **(E)** Quantification of CHI3L3⁺ cells within PDPN-rich areas, demonstrating significantly elevated numbers in EAE versus healthy controls. Data are represented as mean ± SEM. n = 3 per group **(F-G): (F)** Representative images of ARG1⁺ (green) immunoregulatory macrophages in healthy and EAE 3.0 mice, showing spatial colocalization with PDPN⁺ (red) and LYVE-1⁺ structures (white) near olfactory nerves. Insets highlight increased density and clustering of ARG1⁺ cells in EAE. **(G)** Quantification of ARG1⁺ macrophages in PDPN⁺ regions, showing a significant increase in EAE mice compared to healthy controls (p = 0.0116). Data are represented as mean ± SEM. n=6-9 **(H)** High-resolution confocal image of EAE 3.0 olfactory nerve region showing ARG1⁺ macrophages interacting closely with PDPN⁺ and LYVE-1⁺ lymphatics around olfactory nerve bundles (ON). Individual channels shown for ARG1, PDPN, and LYVE-1. Zoomed panel shows perineural ARG1.

### Characterization of CD11c^+^CD11b^+^ immune cells at the inflamed cribriform plate PME

Leveraging podoplanin as an efficient marker of the non-parenchymal meningeal-lymphatic environment at the cribriform plate (**Figure 1**), and the established affinity of CD11b^+^CD11c^+^ cells to this niche (Hsu et al., 2022), we utilized a doublet sorting approach to enrich cell-cell interactions of interest from inflamed cribriform plate preparations during EAE (Bendall, 2020; Hsu et al., 2022) **(Figure 3A**). Specifically, we sorted for CD11b^+^CD11c^+^ cells, which are bound in PDPN^+^CD31^+^ aggregates (**Figure 3B; Supplementary Figure 1**). We also sorted for singlet CD11c^+^/CD11b^+^/CD45^+^cells and singlet cribriform plate PDPN^+^/ CD31^+^/CD45_low_ populations for comparison into a second tube (**Figure 3B; Supplementary Figure 1**). This procedure yielded a tube of sorted singlet no-contact cells and another tube of aggregating cells in heterotypic interactions. Both tubes were then put through a dissociation procedure using Liberase TL (Roche), which yielded post-contact singlets from the cell aggregates for traditional scRNAseq processing (**Figure 3B; Supplementary Figure 1**). This method is distinct from physically-interacting cell sequencing (PIC-seq) (Giladi et al., 2020), whereas in PIC-seq, doublet cells are sequenced in active binding events, and then post-sequencing algorithms are used to deconvolute expression data back to single cell resolution. Here, with PostContact-seq the pipeline is more simplified, and in our experiment involves the isolation of newly dissociated myeloid cells which were previously in contact with a PDPN+CD31+ aggregates without the need for deconvolution algorithms. In essence, this process allows us to interrogate the expressional profile of cells with a previously defined affinity for each other.

**Figure 3.**
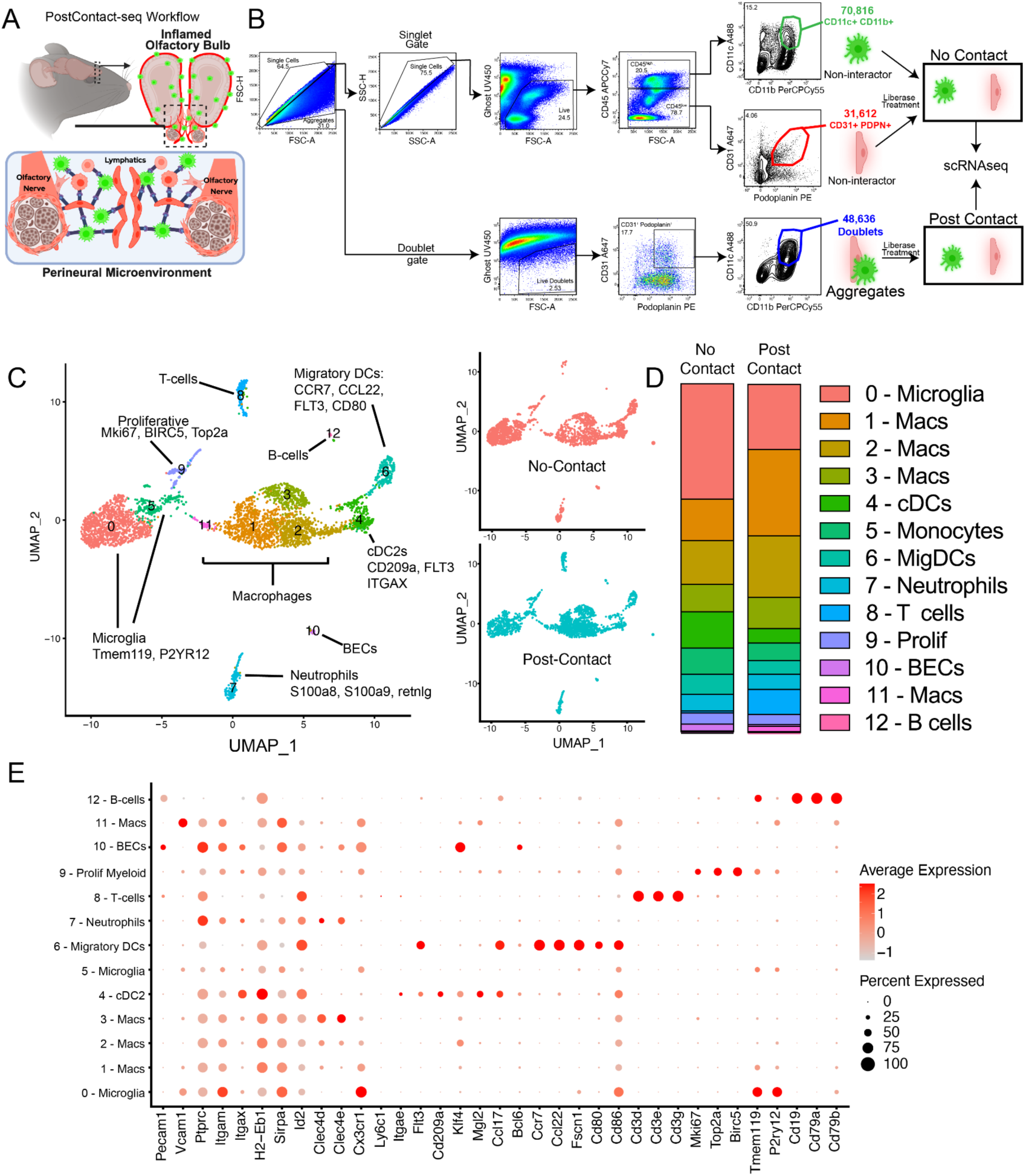
Characterization of post-contact CD11c^+^CD11b^+^ cells which previously interacted with cribriform plate meningeal-lymphatic niche. **(A-B):** Cartoon scheme outlining “PostContact-seq” sorting procedure of cribriform plate + Olfactory bulb tissue preparations from EAE 3.0. Sorted singlets from CD11c^+^CD11b^+^ (Myeloid cells) and CD31^+^PDPN^+^ (Meningeal-Lymphatic niche) gates were collected into a no-contact tube. Simultaneously, quadruple positive CD11c^+^CD11b^+^CD31^+^PDPN^+^ (Myeloid cell+Meningeal Lymphatic Niche) aggregating cells were sorted into the “interactor” tube. Prior to scRNAseq, both tubes underwent short liberase treatment to dissociate interacting cells and generate single cell suspensions of post-contact, and sequenced. **(C):** Combined UMAP plots show the 12 cluster identities from post-contact and no-contact scRNAseq (Left). Overlaid UMAP plots show the 12 cluster identities from post-contact and non-contact scRNAseq (Right). **(D):** Vertical slice graph shows proportion of each cell cluster between no contact and post-contact tubes. **(E):** Dot plot displays the expression of selected marker genes across cell clusters identified by scRNA sequencing.

UMAP analysis revealed 12 distinct clusters (**Figure 3C)**. Overall, no-contact and post-contact clusters had several overlapping cluster classifications (**Figure 3C**), but had fluctuations in the proportion of immune cells in each cluster (**Figure 3D**). The heterogeneity of myeloid cells reflects the fact that CD11b and CD11c are present on both resident and infiltrating macrophages in the CNS during neuroinflammation (Jordão et al., 2019; Wlodarczyk et al., 2014; Mrdjen et al., 2018). Notably, the post-contact sample had elevated levels of three macrophage clusters (Mac 1-3), which were characterized by high *Arg1* expression and other commonly reported tissue-repair, remodeling, and debris clearance signatures (*Chil3, Spp1, Mrc1*, *Apoe, Fn1, Tgfbi)*. Additionally, while the no-contact sample was largely devoid of a T-cell cluster (Cluster 8), the post-contact sample saw a ∼13x fold increase in T-cells (**Figure 3D**). We hypothesize that this could be due to T-cells “hitchhiking” in our sorted aggregates due to their binding affinity to both cell types. While inclusion of post-contact T-cells in the sample preparation was unintended, sequencing revealed aspects of a classical Th1 signature with expression of *Cd3e*, *Cd4, Ifng, Il2rb.* Of note, one of the most prominent clusters, Cluster 0, expressed CD11c (*Itgax*), but also genes related to CNS resident microglia (*Tmem119*, *P2ry12, Siglec-H*) and not the traditional expression signature of dendritic cells (**Figure 3E**). The presence of microglia in the sorted sample supports the presence of CD11c^+^ microglia in the CNS during inflammatory disease (Keren-Shaul et al., 2017), which were included in our CD11c^+^ CD11b^+^ gating strategy. However, while there was a nearly 50% reduction of this microglia population in the post-contact group (**Figure 3D**), some of this sorted population could also be derived from nerve bundle resident macrophages, which resemble microglia (Wang et al., 2020).

Additionally, while our results revealed a more heterogeneous CD11b^+^CD11c^+^ population than anticipated, our scRNAseq analysis revealed two distinct dendritic cell clusters, which correspond to the traditional expressional signatures of cDCs (Cluster 4: *cD209a*, *Flt3*) and Migratory DCs (Cluster 6: *Ccr7*, *Pdcd1lg2, Cd80, Cd83, Cd200*) (**Figure 3E**). Migratory DCs expressed several noticeable Th2 response genes, *IL4I1*, *IL4ra*, *Ccl17*, *Ccl22* and *Bcl2l1* (**Figure 3E**). Broadly, the gene signature of these CCR7^+^ migratory DCs highly overlapped with published scRNAseq datasets of dendritic cells with a “mregDCs” cell state (or “mature DCs enriched in immunoregulatory molecules”) (Maier et al., 2020). This mregDC identity is indicative of a cell state associated with both antigen uptake and maturation but also a restrained stimulatory ability which can be acquired by multiple DC subsets (Maier et al., 2020; Ginhoux et al., 2022).

### Analysis of cribriform myeloid cells supports immunoregulatory retainment at inflamed cribriform plate

To understand the retention of immune cells in the PDPN^+^ PME, we next investigated specifically the post-contact cells from the scRNAseq dataset for clusters that had measurable expression of *Pdpn*. Again, post-contact which we use here refers to previously PDPN-attached sorted cell aggregates which were dissociated prior for scRNAseq. Interestingly, a subpopulation of fibroblasts that expressed the highest levels of *Ppdn* along with several collagen subunits of the ECM: *Col1a1*, *Col1a2*, *Col6a1*, and *Col6a2* (**Figure 4A**). To understand what interactions were implicated in our sorted aggregates, we utilized TALKien, a cell interaction analysis to understand the broad profile of cell-cell crosstalk among cells in the post-contact clusters (**Figure 4B**) (Moratalla-Navarro et al., 2023). Importantly, this integrated analysis utilizes Ligand-Receptor databases from CellChat, CellPhoneDB, iCellNet, and Ramilowsky datasets (Moratalla-Navarro et al., 2023). Cell interaction analysis predicted high levels of crosstalk between macrophages-fibroblasts and macrophages-migratory DC subsets (**Figure 4B-C**). Importantly, this analysis agrees with our immunohistochemistry analysis of the inflamed cribriform plate environment that displays a dense PDPN^+^ meningeal lymphatic environment with interacting myeloid cells (**Figure 4D**, **Figure 1B-J**). Gene ontology analysis of potential interactions pointed to several ECM pathways, but also IL-10 signaling pathways (**Figure 4E**).

**Figure 4.**
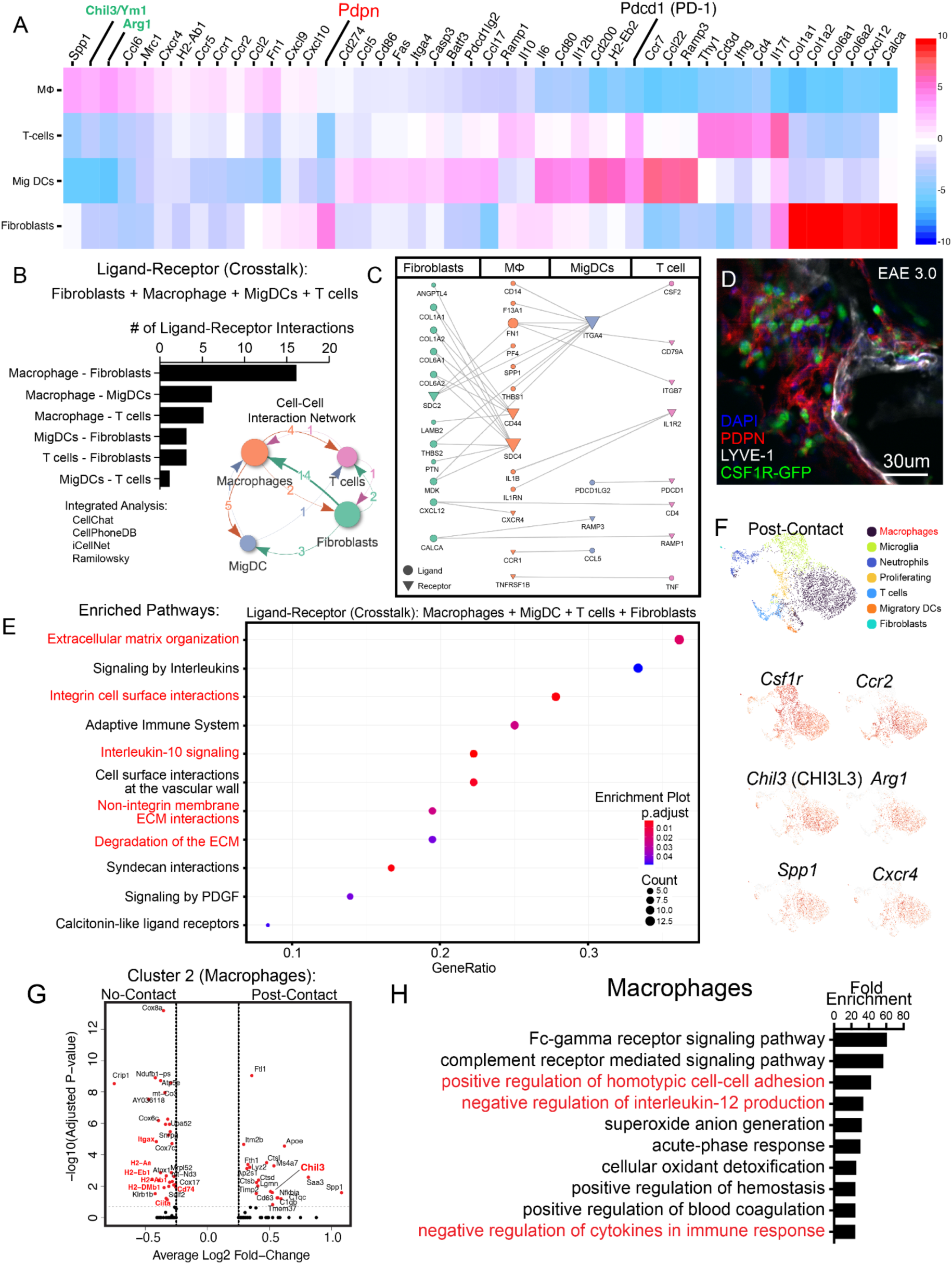
Analysis of post-contact cells reveals immune cell and extracellular matrix crosstalk. **(A)** Heatmap displaying scaled expression (z-score) of highly expressed representative genes involved in immune regulation, immune activation (e.g., *Pdcd1*) across four cell types: macrophages (MΦ), T cells, migratory dendritic cells (Mig DCs), and fibroblasts (e.g., *Pdpn*). Color scale indicates expression levels. **(B)** Quantification and network map of predicted ligand-receptor interactions between fibroblasts and immune populations (macrophages, Mig DCs, and T cells) using integrated analysis from CellChat, CellPhoneDB, iCellNet, and the Ramilowsky dataset. The bar graph shows the number of unique ligand-receptor interactions per pairwise combination, with the most extensive communication observed between macrophages and fibroblasts. Analyzed with crossTALK IntEraction Network (TALKIEN) (Moratalla-Navarro et al., 2023) with microglia, neutrophils, and proliferating gene sets excluded. **(C)** Network plot illustrating specific ligand-receptor pairs involved in fibroblast-immune cell communication. Arrows point from ligand-expressing to receptor-expressing cell types. Notable interactions include *Spp1-CD44*, *Pdpn-Clec2d*, and *Thbs1-CD47*. **(D)** Immunofluorescence imaging of the cribriform plate region in EAE day 3.0 mice showing spatial organization of LYVE-1^+^ lymphatics (white), PDPN^+^ lymphatics and ECM (red), and CSF1R-GFP^+^ myeloid cells (green). DAPI (blue) marks nuclei. Scale bars: 30 µm. **(E)** Pathway enrichment analysis of ligand-receptor pairs among macrophages, Mig DCs, T cells, and fibroblasts. Enriched pathways include extracellular matrix organization, integrin-mediated interactions, interleukin 10 signaling. Gene ratio represents the proportion of identified genes contributing to each pathway; dot size reflects the number of genes, and color indicates adjusted p-value. **(F) :** UMAP projection shows major post contact cell types, including macrophages (purple), microglia (green), neutrophils (blue), T cells (orange), proliferating cells (light blue), migratory dendritic cells (yellow-orange), and fibroblasts (aqua). Feature plots highlight expression of *Chil3* (CHI3L3), *Arg1*, *Spp1*, *Ccr2*, *Cxcr4*, *Csf1r* **(G) :** Volcano plot comparing post-contact vs no-contact cluster 2 macrophages (Highest *Arg1* expressing cluster). Analysis reveals post contact macrophage have downregulated MHC-II processing and presentation genes (*H2-DMa*, *H2-Aa*, *H2-Ab1*, *H2-DMb1*, *H2-Eb1*, *Cd74*, *Ciita*) and elevated expression of *Chil3* (CHI3L3), another alternatively activated macrophage marker and ECM binding protein. **(H) :** Gene ontology enrichment analysis (bottom) demonstrates post contact Macrophages are enriched for pathways in Fcγ receptor and complement receptor signaling, negative regulation of IL-12 production, homotypic cell-cell adhesion, and hemostasis.

Additionally, Th2-skewed macrophages are characterized by high expression *Arg1*^+^/*Chil3*^+^ and are the dominant post-contact cluster in our dataset (**Figure 4F, Supplemental Figure 3**) and previously found to increase at cribriform plate (**Figure 2**). Interestingly, post-contact macrophages (Cluster 2) have lower expression of MHC-II antigen processing and presentation genes (*H2-DMa*, *H2-Aa*, *H2-Ab1*, *H2-DMb1*, *H2-Eb1*, *Cd74*, *Ciita*) and elevated expression of *Chil3*, *Sdc4*, and *Spp1* key ECM related binding proteins (**Figure 4G**). This population also expressed *Ccr2* and *Cxcr4* (**Figure 4F**), implicating monocyte recruitment to cribriform plate may occur via classical CCL2-CCR2 and CXCR4-CXCL12 axis to the ECM meningeal regions. In agreement with this CCR2^+^ cells increased significantly in PDPN^+^ regions between olfactory bulbs during EAE, even appearing to embed into a PDPN^+^ ECM (**Supplemental Figure 4A-C**). In agreement with an alternatively activated phenotype, enriched gene ontology pathways for post-contact macrophages included homotypic cell adhesion (*Pdpn*, *Emilin2*, *Lgals1*, *Plaur*), ECM-formation (*Ecm1, Tgfbi*, *Emilin2*, *Fn1),* negative regulation of IL-12 production, and negative regulation of cytokines in immune response (**Figure 4H**).

### Cribriform DCs have enrichment pathways related to reduced pro-inflammatory signature and *Pdcd1* (PD-1) expression

CCR7^+^ migratory DCs (cluster 6) were a key post-contact subset in our data set, and MHCII^+^ cells were heavily implicated in our IHC analysis (**Figure 5A**). Enriched pathways in cluster 6 DC included positive regulation of cell adhesion, negative regulation of immune system processes, and positive regulation of programmed cell death (**Figure 5B**). Overall, post-contact and no-contact DCs shared a highly overlapping signature, but further comparison of enrichment pathways from the top 300 gene list of no-contact and post-contact DCs revealed several unique pathways within the post-contact DCs including, programmed cell death, caspase activation, tolerance induction, and response to L-arginine (**Figure 5C**). Inversely, we saw several unique pro-inflammatory gene enrichment pathways in the no-contact DCs, including positive regulation of IL12 and IL17 signaling (**Figure 5C**).

**Figure 5.**
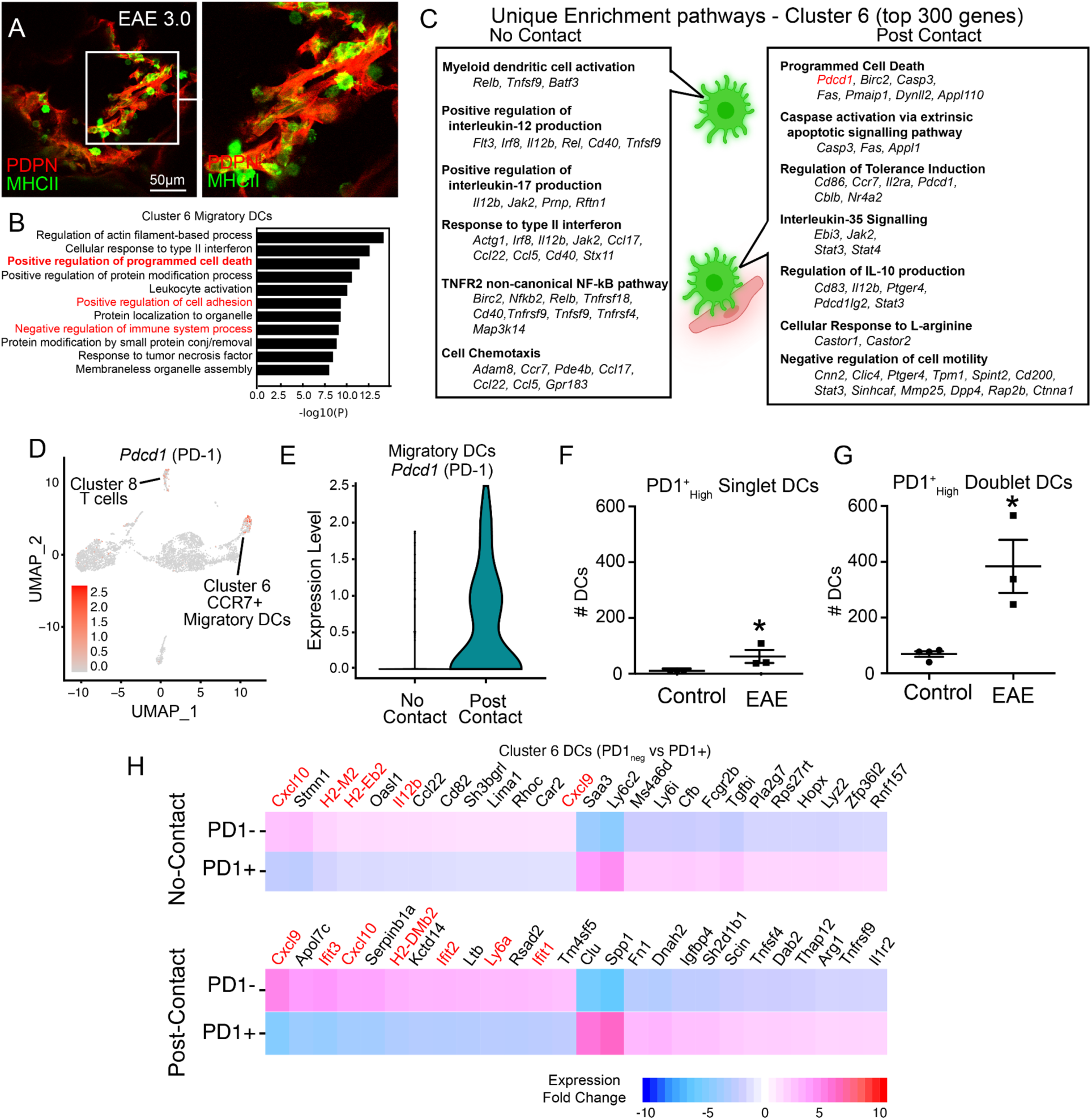
Post-Contact DCs have pathways related to reduced pro-inflammatory signature and *Pdcd1* (PD-1) expression. **(A)** High-resolution confocal image from EAE 3.0 CP showing MHCII⁺ (green) cells in close proximity to PDPN+ stroma (red). **(B)** Gene ontology enrichment analysis of cluster 6 migratory DCs reveals pathways related to actin dynamics, interferon signaling, leukocyte activation, and programmed cell death. Red text indicates immune pathways of interest. **(C)** Comparison of unique enrichment pathways from top 300 genes compared between no-contact and post-contact DCs. **(D)** UMAP visualization of scRNA-seq data highlighting *Pdcd1* (PD-1) expression in cluster 6 CCR7⁺ migratory DCs and cluster 8 T cells **(E)** Violin plot showing *Pdcd1* expression in post contact DCs versus no-contact, suggesting PME influence on DC checkpoint expression. **(F-G)** Quantification of PD-1^High^ single and doublet DCs at the olfactory-cribriform junction, showing significant increase in EAE mice compared to controls. n = 3, unpaired Student’s t-test. Data are represented as mean ± SEM. Singlet DCs (p=0.0322), Doublet DCs (p=0.0084). **(H)** Heatmaps of highest genes expressed genes in PD-1⁺ vs PD-1⁻ DCs within CD31+PDPN+ post-contact and no-contact subsets. Pro-inflammatory genes (*Cxcl10*, *Il1b*, *Ifit1, Ifit2, Ifit3*.) are high in PD-1 negative DCs.

Focusing on programmed cell death pathways, we next sought to identify all the immune cells within our scRNAseq dataset that express *Pdcd1*, the gene which encodes programmed cell death receptor 1 (PD-1), a well-characterized checkpoint inhibitor receptor expressed in activated infiltrating CD4 T-cells during EAE (Schreiner et al., 2008). While *Pdcd1* was expressed in the post-contact T-cell cluster (Cluster 8), it was also detectable in the CCR7+ migratory DCs (Cluster 6) (**Figure 5D**). Furthermore, when we further divided the data into “post-contact” vs “no-contact” clusters, we noticed that *Pdcd1* was primarily expressed by the post-contact migratory DCs (**Figure 5E**). Using a minimum threshold of 0.5 *Pdcd1* expression level, we estimated that around 45.8% of the post-contact migratory DCs were positive for *Pdcd1* expression, whereas <15% of no-contact migratory DCs were PD-1^+^. To verify surface level expression of PD-1 on cribriform migratory DCs during neuroinflammation, we next performed flow cytometry analysis on leukocytes in cribriform plate cell suspensions during EAE and steady state. Here, similar to the scRNAseq experiment, we gated for singlet and doublet CD11c^+^ CD11b^+^ CD45^+^ DCs and identified that singlet DCs in our cribriform plate preparation contain PD-1 at detectable levels during EAE, and that doublet DC aggregates expressed PD-1 (**Figure 5F-G, Supplemental Figure 2)**.

Engagement of PD-1 on myeloid cells, including DCs, has been shown to inhibit interferon (IFN) signaling pathways through the activation of SHP-2 (Knutson, 2024; Klement et al., 2023). This inhibition in DCs has been shown to lead to reduced production of pro-inflammatory chemokines and cytokines mainly: CXCL9, CXCL10, and IL12B, which are crucial for pro-inflammatory T cell recruitment and initiating Th1 immunity by DCs (Knutson, 2024; Klement et al., 2023). To better understand the inflammatory profile of PD-1^+^ DCs at the cribriform plate, we compared the expression signature of PD-1 expressing and non-expressing CCR7^+^ DCs from cluster 6 of our datasets (**Figure 5H**). Highest genes in PD1^negative^ DCs included chemokines *Cxcl9*, *Cxcl10*, *Il12b*, interferon-Stimulated Genes (*Ifit1*, *Ifit2*, *Ifit3*), and several MHC-related genes (*H2-M2*, *H2-Eb2*, *H2-DMb2)* (**Figure 5H**). Together, these data support that migratory DCs expressing *Pdcd1* exhibit an immunosuppressive gene signature which has been recently described in myeloid cells, including DCs, in tumor microenvironments (Knutson, 2024; Klement et al., 2023). Our data extends these described characteristics of PD-1+ DCs to the cribriform plate drainage environment during EAE-induced autoimmunity.

### Peripheral blood derived DCs undergo cell death at cribriform plate-lymphatic niche during neuroinflammation

Since some DCs displayed some unique pathways related to programmed cell death and apoptosis, we next sought to better understand the kinetics of recruitment and death of CD11c^+^CD11b^+^ cells at the cribriform plate lymphatic niche during EAE. To do this, we utilized a method of CD45 intravascular staining (CD45-IV) to tag blood-originating leukocytes and track their migration into the olfactory-cribriform plate axis. This IV-staining method originally, established by *Anderson et al.,* can label ∼99% of leukocytes in the blood within 3 minutes of injection; these labeled leukocytes can then be tracked out of the blood and quantified in tissues across time (Anderson et al., 2014). Using a retro-orbital injection, we IV-stained peripheral blood CD45^+^ immune cells at several time points prior to tissue collection (3 minute, 6, 9,12, 18, 24, 48 hrs). Then we investigated the presence of CD45 IV^+^ DCs bound to CD31^+^PDPN^+^ aggregates at the cribriform plate (**Figure 6A; Supplementary Figure 5**). Here, we found that at the 3-minute time point, very few of the doublets (<1%) originated directly from the blood in our sample preparation (**Figure 6A)**, indicating that there is minimal contamination from blood. Interestingly, between 24-48 hours after IV-injection, the total percentage of CD45IV^+^ stained DCs increases steadily (**Figure 6B**), highlighting a time-dependent relationship to CD11c^+^CD11b^+^PDPN^+^CD31^+^ binding events. We also determined that doublets had increasing Ghost^+^ cells across time, which suggests cells are dying in a time-dependent relationship (**Figure 6C**). To validate the apoptotic potential of DCs at the cribriform plate, we stained EAE sections from CD11c-eYFP mice with Cleaved Caspase-3 and Lyve-1 (**Figure 6D**), where we confirmed Cleaved-Caspase-3^+^ DCs in an active binding interaction with Lyve-1 (**Figure 6D**).

**Figure 6.**
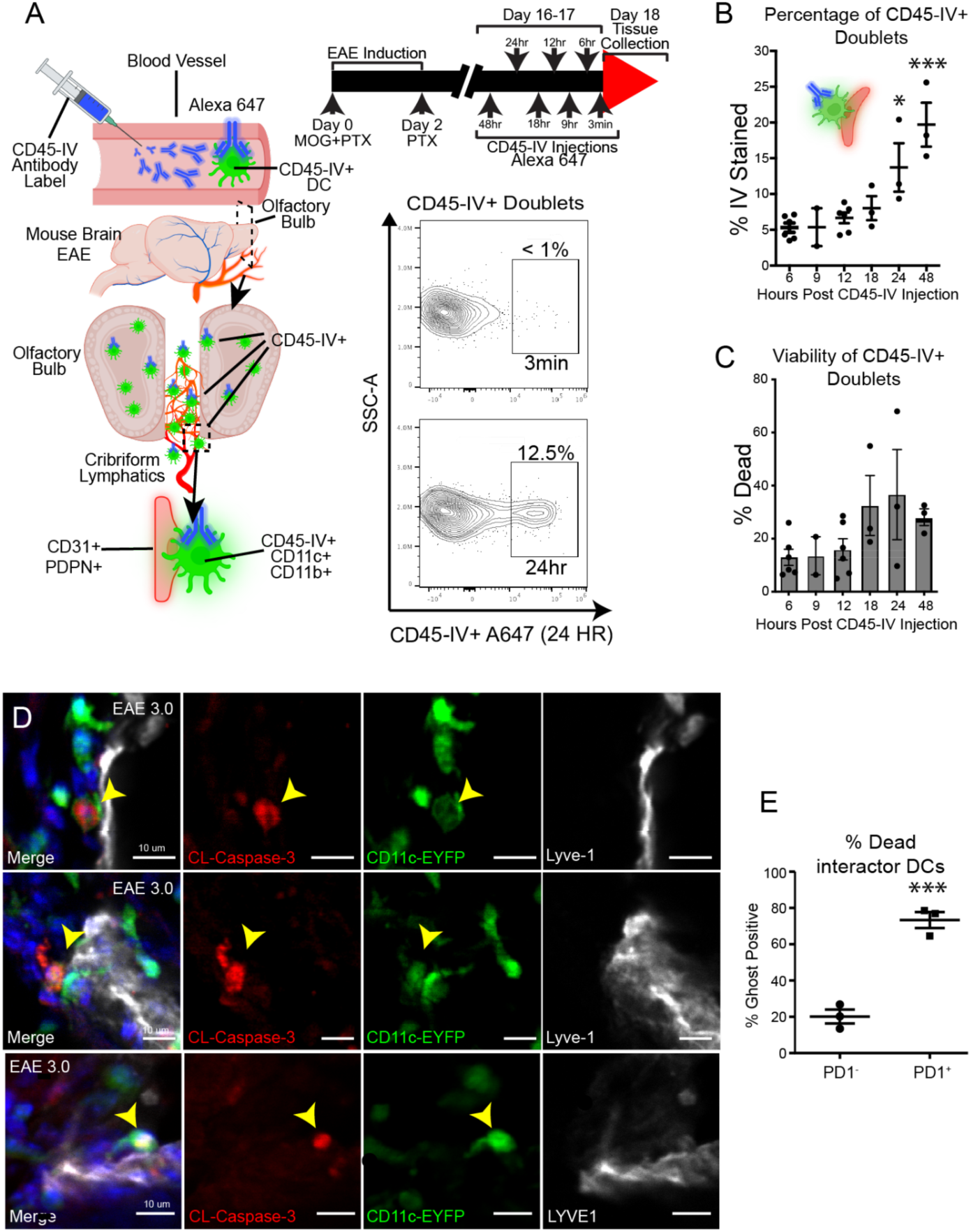
Blood-derived CD11c+CD11b+ cells migrate to cribriform plate lymphatics in a time dependent manner and undergo cell death. **(A)** Schematic overview of experimental design. Mice were intravenously injected with Alexa Fluor 647-labeled CD45 antibody (CD45-IV) at multiple timepoints following EAE induction to label blood-accessible leukocytes. Diagram (left) shows DC trafficking to the cribriform plate niche. Representative flow plots (right) show the lack of CD45-IV⁺ doublets at 3 minutes and emergence of IV+ cells at 24 hours post-injection. **(B)** Quantification of the percentage of CD45-IV⁺ CD11c+CD11b+CD31+PDPN+ doublets over time, showing progressive accumulation of stained doublets, peaking at 48 hours post-injection. 24 hr (p=0.0227) and 48 hr (p=0.0001) time points are significantly elevated when compared 6 hr. One way anova with Tukey’s multiple comparison. Data combined from 3 independent experiments. Data are represented as mean ± SEM n=2-6 per group **(C)** Viability of CD45-IV⁺ doublets assessed by live/dead staining at different timepoints, showing trend of cell death (Ghost+) over time. One-way anova (P = ns). data are represented as mean ± SEM n=2-6 per group **(D)** Representative confocal images from EAE 3.0 mice showing cleaved caspase-3⁺ (red) CD11c-EYFP⁺ DCs (green) interacting with LYVE-1⁺ lymphatics (white). Yellow arrowheads highlight apoptotic DCs. **(E)** Quantification of Ghost+ cells among PD-1⁻ and PD-1⁺ DCs identified as interactors, showing significantly higher cell death among PD-1⁺ interactor doublets. n = 3, P = 0.0008, unpaired Student’s t-test. Data are represented as mean ± SEM.

Finally, we analyzed the viability of PD-1^+^ DC doublet aggregates using flow. We determined that PD-1^+^ aggregates had a higher percentage of Ghost^+^ cells when compared PD1^neg^ DC doublet aggregates, suggesting PD-1^+^ DCs undergo cell death in engagement with this niche (**Figure 6E)**. Together, these data support that blood-derived CD11c+ DCs can migrate to the cribriform plate PME regions in a time-dependent manner, with subpopulations dying as they interface with the meningeal-lymphatic environment.

## Discussion

Drainage of cerebrospinal fluid (CSF) and its associated inflammatory cues are essential for maintaining CNS fluid homeostasis, clearing waste, and coordinating neuroimmune responses with the periphery (Kipnis, 2024). Recent work by our lab and others has demonstrated that lymphatic vessels associate with CSF-interfacing olfactory nerve bundles at the cribriform plate and undergo dynamic remodeling during neuroinflammation, where they undergo lymphangiogenesis, facilitate antigen drainage, and upregulate transcriptional programs linked to leukocyte cross-talk (Hsu et al., 2022, 2019; Spera et al., 2023; Decker et al., 2022; Xin et al., 2024). Previously, we determined that the cribriform plate niche is dominated by interacting CD11c⁺CD11b⁺ myeloid cells (Hsu et al., 2022) with access to nearby skull bone marrow via perineural bone channels (Laaker et al., 2025). Together, these findings support the concept that the perineural microenvironment (or PME) at the cribriform plate functions as a specialized immunoregulatory interface, linking the previously described CSF-to-lymphatic drainage with the recruitment, retention, and programming of PME myeloid cells.

In this study, we isolated CD11c^+^CD11b^+^ cells in contact with PDPN^+^CD31^+^ aggregates from inflamed cribriform plate preparations, dissociated cells from their ongoing cell-cell interactions, and then analyzed their residual post-contact gene expression profile compared to non-interacting controls. Our analysis revealed several immune cell interactions but suggested a high number of macrophage, migratory DC, and fibroblast (ECM) interactions. Additionally, no-contact DCs had unique pathways more characteristic of activated DCs, including IL-12, IL-17 production, and levels of chemokine receptors. In contrast the post-contact migratory DCs were enriched with pathways related to immune tolerance, reduced cell motility, and programmed cell death including the expression of the inhibitory receptor *Pdcd1* (PD-1). In summary, retained DCs had more regulatory and sessile characteristics from our analysis.

While many potential interaction pathways emerged, we decided to interrogate programmed cell death and Th2-skewed immune responses at the cribriform plate. Interestingly, in EAE, PD-1^+^ DCs accumulated at higher levels at the cribriform plate region, and PD-1 expression had a signature associated with inhibited interferon signaling. Immunosuppressive signatures extended to macrophage subsets too, with elevated ARG1^+^/CHI3L3^+^ alternatively activated macrophages accumulating in these PDPN^+^ regions during EAE. Together, our findings support the establishment of an immunosuppressive myeloid network in the PDPN^+^ ECM surrounding CSF-effluxing and lymphatic interfacing olfactory nerve bundles (**Supplemental Figure 6)**.

Previously, our lab showed that *Cd274* (PD-L1), the ligand for PD-1, increases in the cribriform plate lymphatic niche during EAE, suggesting neuroinflammation promotes suppressive PD-L1/PD-1 crosstalk at this site (Hsu et al., 2022). Additionally, Pdcd1lg2 (PD-L2), Cd80, and Cd83 by migratory DCs likely reflects a tolerogenic activation state rather than a conventional immunostimulatory. While PD-1’s inhibitory role on effector responses in T-cell subsets has been extensively studied, its role in DCs is considerably less understood, though documented in several disease contexts. For example, PD-1^+^ DCs have been identified in the tumor microenvironment (TME) in both human and animal studies (Karyampudi et al., 2016; Strauss et al., 2020; Kwiecień et al., 2022; Lim et al., 2016; Krempski et al., 2011; Wang et al., 2022b; Gordon et al., 2017). Similar to its function in T-cells, PD-1 on DCs has been implicated as highly inhibitory, reducing DC stimulatory ability and survival, and negatively influencing tumor response (Karyampudi et al., 2016). Inversely, eliminating PD-1 in *Pdcd1* KO, APCs exhibit an increased ability to activate T-cells in a tumor model, resulting in better tumor control (Strauss et al., 2020). Interestingly, Park and colleagues showed that *Pdcd1*-KO DCs were less likely to be apoptotic and were able to evoke higher antigen-specific T cell responses (Park et al., 2014). Thus PD-1 appears to be a useful receptor to identify subsets of inhibited DCs. In agreement with this, our data support that *Pdcd1*-expressing DCs also have low levels of *Cxcl9, Cxcl10, Il12b,* and several antigen presentation and processing genes. PD-1 engagement by PD-L1 has been proposed to suppress IFN-signaling pathways in DCs through the SHP-2 pathway (Knutson, 2024; Lamichhane et al., 2017). In line with this, several Interferon-Stimulated Genes (*Ifit1*, *Ifit2*, *Ifit3*) were lower in PD-1^+^ DCs in the cribriform plate dataset.

While the induction pathways of *Pdcd1* in DCs is not fully elucidated, evidence suggests that sustained IL-10 secretion from nearby myeloid cells (Macrophages and/or MDSCs) is a key pathway that upregulates the PD-1 receptor on DCs (Knutson, 2024; Lamichhane et al., 2017). Notably, IL-10 secretion is commonly associated with ARG1+ / CHI3L3+ producing macrophage subsets, which we identified both in our scRNAseq dataset and in IHC analysis of PME-interacting cells. Additionally, our data supports that Arg1^+^ macrophages express PDPN as well (Zhang et al., 2025a; Wu et al., 2024), contributing to niche complexity. Analysis suggests that IL-10 and ECM remodeling pathways are enriched in the immune-stromal environment surrounding olfactory nerves. In models of sciatic nerve injury, infiltrating monocyte-derived macrophages are recruited and retained perineurally, adopting an *Arg1^+^* / *Chil3^+^*tissue repair-associated program, which is distinct from resident macrophage subsets (Ydens et al., 2020). Thus, our observation of a similar type 2 skewed immunoregulatory niche around olfactory nerve bundles may also promote local tissue remodeling and immunosuppression at the cribriform plate PME, a known site of CSF and antigen drainage(Hsu et al., 2022; Laaker et al., 2025).

In a previous study, we identified in mice that CSF-interacting olfactory nerve bundles have connections to a nearby pool of ethmoid bone marrow (Laaker et al., 2025). During EAE, CSF1R-GFP^+^ immune cells could be observed traversing these skull channels, providing a potential pathway for local recruitment and coordination of myeloid immune responses to the PME (Laaker et al., 2025). As a result, CCR2+ monocytes and CCR7+ DCs may migrate or mature at the cribriform plate lymphatic region and adopt these tolerogenic characteristics.

Similar immunosuppressive stromal-lymphatic environments are also commonly reported in tumor microenvironment (TME), and are known to promote suppressive macrophages and tolerogenic DCs (Swartz and Lund, 2012; Bieniasz-Krzywiec et al., 2019; Xiao et al., 2023). Additionally, local metabolic shifts in the TME have been implicated as a driver of tolerogenic DC cell states (Snyder and Amiel, 2018). Direct depletion of nutrients such as arginine by M2 macrophages and MDSCs can lead to mTOR inactivation in DCs, which directly reduces DC stimulatory ability (Turnquist et al., 2007; Sukhbaatar et al., 2016; Reichardt et al., 2008; Mondanelli et al., 2017). Future studies will investigate which factors can most reliably influence a broad immunosuppressive environment surrounding CSF-nerve drainage sites. These factors include TGF-β, which is also secreted by alternatively activated macrophages and has been shown to highly induce meningeal stroma thickening and suppress antigen outflow from meningeal lymphatics (Hitpass Romero et al., 2025).

There are several limitations to our investigation. For one, our sequencing approach prioritized myeloid cell-cell interactions, and reduced the acquisition of important stromal and endothelial cell populations in downstream sequencing. Additionally, due to anatomical constraints of the region, we did not perform any selective targeting or depletion of these myeloid cell populations. Future investigations will attempt to selectively inhibit the cribriform myeloid populations to alter disease pathogenesis and utilize complementary approaches like PIC-seq to compare methodology. In addition, we opted to look at a single timepoint of neuroinflammation at peak EAE.

In summary, our findings support that the cribriform plate lymphatic region is a dynamic site of myeloid cell regulation, which is enriched for Th2-skewed macrophages and a subpopulation of migratory DCs with a tolerogenic signature during EAE. Given cribriform lymphatic vasculature facilitates DC drainage to cervical lymph nodes (Hsu et al., 2019, 2022; Jin et al., 2025; Yoon et al., 2024), these findings could be important for altering DC phenotypes and coordinating peripheral immune responses. Our data also highlights the complexity of the cribriform plate region, with the integration of anti-inflammatory macrophages, which likely play a role in protecting local nerve health, promoting tolerance and modulating lymphatic outflow of region (Hsu et al., 2022; Choi et al., 2025; Hsu et al., 2019). These results raise the question of if targeted delivery of immunotherapies at DC efflux sites and immune hub regions could promote more efficent CNS immune response during diseases like brain cancer, Alzheimer’s, or CNS infection (Li et al., 2023). Inversely, enhancing the immunosuppressive capabilities at skull foramina sites may be a new aspect of treating CNS autoimmunity. However, comparable anatomical and immunoregulatory relationships need to be fully investigated in humans. Future studies will attempt to test this framework to understand how drainage sites can tune APCs for downstream impacts on peripheral immunity and further elucidate the unique features of these environments (Hsu et al., 2022; Laaker et al., 2025).

## Materials and Methods

### Animals

Female C57BL/6J wild-type (stock #: 000664) and CSF1R-GFP MacGreen mice (stock #: 018549), were purchased from Jackson Laboratories. CD11c-eYFP transgenic reporter mice were a generous gift from Dr. Michel C. Nussenzweig at Rockefeller University. Eight to twelve-week old female mice were used for all EAE experiments. All experiments were conducted in accordance with guidelines from the National Institutes of Health and the University of Wisconsin, Madison Institutional Animal Care and Use Committee.

### EAE Induction

EAE was induced in 8 to 12-week-old female mice by subcutaneous immunization with 100 µg of MOG_35-55_ emulsified in Complete Freund’s Adjuvant (CFA) between the shoulder blades. 200 ng of Pertussis Toxin (PTX) was injected intraperitoneally at 0 days post immunization (d.p.i.) and 2 d.p.i. The onset of clinical scores were observed between day 8 and 12 post-immunization, and were assessed daily as follows: 0, no clinical symptoms; 1, limp/flaccid tail; 2, partial hind limb paralysis; 3, complete hind limb paralysis; 4, quadriplegia; 5, moribund. Intermediate scores were also given for the appropriate symptoms. An EAE clinical score between 2.5-3.5 at day 15 to 18 post-immunization was used for all experiments.

### Histology

Mice were terminally anesthetized with isoflurane and transcardially perfused with 0.1M PBS followed by perfusion with 4% paraformaldehyde (PFA) in 0.1M PBS. Mice were then decapitated and the skin surrounding the head was removed using forceps and scissors to separate the skin from the muscle and ear canal. The whole heads were fixed in 4% PFA in 0.1M PBS overnight. The whole heads were then decalcified in 14% ethylenediaminetetraacetic acid (EDTA) in 0.1M PBS for 7 days followed by cryoprotection in 30% sucrose in 0.1M PBS for 3 days. The EDTA was replaced with fresh 14% EDTA each day. The decalcified mouse heads were then embedded in Tissue-Tek OCT Compound, frozen on dry ice, then stored at -80°C. 60 𝜇m thick frozen sections were obtained on a Leica CM1800 cryostat, mounted on Superfrost Plus microscope slides and stored at -80°C. For dural tissues, the skullcap was isolated and collected after decalcification and stored at 4°C in a 48-well non-tissue culture treated plate containing PBS.

### Immunohistochemistry

Sections were thawed at room temperature for 10 minutes, washed with 0.1M PBS for 10 minutes, then unspecific binding blocked with 10% bovine serum albumin (BSA) with 0.1% Triton-X for permeabilization in 0.1M PBS for 60 minutes. Sections were then incubated with the appropriate primary antibodies in 1% BSA and 0.1% Triton-X in 0.1M PBS at 4°C overnight in a humidified chamber. The following antibodies were used for immunohistochemistry: Podoplanin PE (eBioscience, Catalog #: 12-5381-80), CD31 Alexa647 (BD Biosciences, Catalog #: 563608), Lyve-1 eFluor660 (Thermo Fisher Scientific, Catalog #: 50-0443-80), MHC II eFluor450 (eBioscience, Catalog #: 48-5321-80), CD11c Alexa488 (Thermo Fisher Scientific, Catalog #: 53-0114-80), YM1 / CHI3L3 (R&D Systems Catalog #: AF2446), Arginase-1 (Invitrogen, Catalog #: PA5-29645), Cleaved Caspase-3 (Abcam, Catalog #: E83-77). Sections were then washed three times with 0.1M PBS for 10 minutes each, then incubated with the appropriate secondary antibodies in 1% BSA and 0.1% Triton-X in 0.1M PBS at room temperature for 120 minutes. The following secondary antibodies were used: Donkey anti-Chicken Alexa488 (Invitrogen, Catalog #: A11039), Donkey anti-Chicken Alexa647 (Invitrogen, Catalog #: A21449), Donkey anti-Goat Alexa568 (Invitrogen, Catalog #: A11057), and/or Donkey anti-rabbit Alexa568 (Invitrogen, Catalog #: A10042). Sections were then washed three times with 0.1M PBS for 10 minutes each, mounted with Prolong Gold mounting medium with DAPI, and images acquired using an inverted Olympus Fluoview FV1200 confocal microscope.

### Single Cell Suspension Preparation

Mice were terminally anesthetized with isoflurane and transcardially perfused with PBS. Mice were then decapitated, and the skin was cut dorsal to the midline of the skullcap rostrally to expose the brain. The skullcap was then removed along with the brain and dura after separation from the olfactory bulbs. The cribriform plate and its associated tissues, which included portions of the olfactory bulbs and the cribriform plate, were dissected out and placed in a 70-micron strainer submerged in RPMI-1640 in a non-tissue culture treated dish. The tissues were then mechanically dissociated by pushing the tissue through the strainer using a syringe plunger. The mechanically dissociated cells were then spun down, washed, and resuspended in FACS buffer (1% Bovine Serum Albumin in 0.1M PBS) for FACS staining.

### Flow Cytometry

After resuspension of mechanically dissociated cells in fluorescence-activated cell sorting (FACS) buffer (pH = 7.4, 0.1M PBS, 1 mM EDTA, 1% BSA), the cells underwent staining with conjugated antibodies/dyes. Conjugated antibodies were diluted 1:200 in FACS buffer, and the following antibodies and dyes were used: Ghost UV450 (Tonbo Biosciences, Catalog #: 13-0868-T500) or Ghost Violet540 (Tonbo Biosciences, Catalog #: 13-0879-T100) to visualize live/dead cells; CD31 Alexa647 (BD Biosciences, Catalog #: 563608), Podoplanin PE (eBioscience, Catalog #: 12-5381-82), CD45 APC-Cy7 (Biolegend, Catalog #: 103116) or CD45 APC eFluor780 (eBioscience, Catalog #: 47-0451-80) to visualize leukocytes; CD11b PerCP-Cy5.5 (Biolegend, Catalog #: 101227) or PE (Invitrogen, Catalog #:12-0112-82), CD11c Alexa488 (Invitrogen, Catalog #: 2284135), or CD11c FITC (Biolegend, Catalog #: 117305) to visualize dendritic cells; PD-1 BV421 (Biolegend, Catalog #: 109121), CD4 A647 (BD Biosciences, Catalog #: 557681). After staining for 30 minutes at 4°C, cells were washed 3 times with FACS buffer and run using Cytek’s 3-laser Northern Lights.

### FACS Sorting

After generating a single cell suspension and staining for cribriform plate with Ghost UV450 (Tonbo Biosciences, Catalog #: 13-0868-T500), CD31 Alexa 647 (BD Biosciences, Catalog #: 565608), Podoplanin PE (eBioscience, Catalog #: 12-5381-82), and CD45 APC-Cy7 (Biolegend, Catalog #: 103116), the cells were sorted with a FACS Aria III with a nozzle size of 130 um at the UW Flow Core satellite facility in the UW Biotechnology Center. Cells were gated for singlets using both FSC and SSC, dead cells excluded by gating for Ghost^negative^ live cells, and were identified using both CD31^+^ Podoplanin^+^ after excluding both CD45^intermediate^ microglia and CD45^+^ leukocytes as previously described(Hu et al., 2020; Herz et al., 2018; Louveau et al., 2015; Da Mesquita et al., 2018). Cells were sorted into either FACS buffer (1% BSA in PBS) for scRNA-seq, or RPMI containing 10% FBS, 2-mercaptoethanol, and penicillin/streptomycin for *in vitro* co-culture experiments.

### scRNA Sequencing

A single cell suspension of the cribriform plate was generated from 8 pooled EAE mice and FACS sorted for singlets, Ghost^-^ live cells, CD45_low_, Podoplanin^+^, and CD31^+^. Sorted cell suspensions were then provided to the Biotechnology Core facility at the University of Wisconsin Madison for single cell sequencing using the 10x Genomics Chromium Single Cell Gene Expression Assay. A total of 70,816 (CD11b^+^,CD11c^+^) and 31,612 (CD31^+^, PDPN^+,^ CD45_low_) singlets were sorted for the control group. From the same sample preparation, 48,636 doublet (CD11b^+^, CD11c^+^, CD31^+^, PDPN^+^) events/cells were sorted provided to the Biotechnology Core for scRNA-seq. Both singlet and doublet tubes underwent treatment with 0.25 mg/mL Liberase TL (Roche) at 37*C on a shaker for 5 minutes. Samples were then washed and resuspended with a FACS buffer (1% BSA + 1x PBS). Cells were loaded onto a Chromium Controller to generate a single cell + barcoded gel-bead emulsion. Libraries were prepped with the 10X Genomics 3’ reagents kit (v3 chemistry). The target recovery rate was 3,000 cells with a targeted read depth of 76,000 per cell. Cells were sequenced on the NovaSeq S1 100-cycle flow cell in collaboration with the University of Wisconsin Biotechnology Center (UWBC) DNA Sequencing Facility. Data were analyzed by the UW Bioinformatics Resource Center. Experimental data were demultiplexed using the Cell Ranger Single Cell Software Suite, mkfastq command wrapped around Illumina’s bcl2fastq. The Miseq balancing run was quality controlled using calculations based on UMI tools (Smith et al., 2017). Sample libraries were balanced for the number of estimated reads per cell and run on an Illumina NovaSeq system. Cell Ranger software was then used to perform demultiplexing, alignment, filtering, barcode counting, UMI counting, and gene expression estimation for each sample according to the 10x Genomics documentation. Barcodes containing unusually high numbers of detected transcripts indicative of a doublet signature were excluded. The resulting data were then analyzed and explored using the Loupe Cell Browser software and the R packages clusterProlifer and/or monacle3 after excluding cells that made it through the FACS-enrichment. In total, 5115 (no-contact) and 4423 (post-contact) cells were identified in the analysis. 9,538 cells sequenced total (combined from no-contact and post-contact runs). Filtered cells for differential analysis using the following criteria: Cells have > 200 genes expressed, Cells have < 3,500 genes expressed, Cells have < 5% mitochondrial genes expressed. Using these criteria, filtered down to 4,132 cells for downstream analyses. 1,971 and 2,161 cells from no-contact run and post-contact run, respectively.

### Gene ontologies and Ligand Receptor analysis

Genes identified to be differentially expressed from each cluster were separately assessed by gene ontological analyses using R package clusterProfiler (Yu et al., 2012). The genes that passed filtering during alignment were used as the background set during over-representation analyses. Significant gene ontological terms were identified using an adjusted P-value < 0.05. Bar plots, gene-concept network plots, and enrichment maps were visualized and generated using clusterProfiler. For receptor ligand interaction analysis we utilized TALKIEN: crossTALK IntEraction Network (Moratalla-Navarro et al., 2023). TALKIEN integrated analysis utilizes Ligand-Receptor DataBases from four sources: CellChat, CellPhoneDB, iCellNet, and Ramilowsky datasets (Moratalla-Navarro et al., 2023).

### *In vivo* CD45 I.V. labeling

EAE was induced in C57BL/6 wild-type mice, and at peak disease score mice were intravenously (I.V.) injected with 2.5 µg of CD45 antibody conjugated to Alexa647 (Biolegend, Catalog #: 109818) in 200 µL of PBS as previously described (Anderson et al., 2014). 3 minutes before harvest to control for *ex vivo* unspecific binding of blood-derived cells to cribriform plate niche. Unperfused mice were used as positive controls to visualize blood-derived binding of leukocytes to cribriform plate niche, where intravenous delivery of CD45 antibody labeled at least 95% of blood-derived leukocytes after 3 minutes. Perfused mice were used for all experiments. Cribriform plates were then processed for flow as described previously, and processed using Cytek’s 3-laser Northern Lights.

### Statistics

When comparing results from two groups, unpaired Student’s t-test was used. When comparing results from three groups, one-way ANOVA using Tukey’s post-hoc multiple comparisons test was used. When comparing results across time, two-way ANOVA using Sidak’s multiple comparisons test was used. All statistical analysis was performed using Graphpad Prism 6.0 software. Data is portrayed as the mean +/- standard error of the mean (S.E.M.), and significance portrayed as: n.s. = non-significant, *p* > 0.05; \**p* < 0.05; \*\**p* < 0.01; \*\*\**p* < 0.001; \*\*\*\**p* < 0.0001

---

**Supplemental Figure 1.**
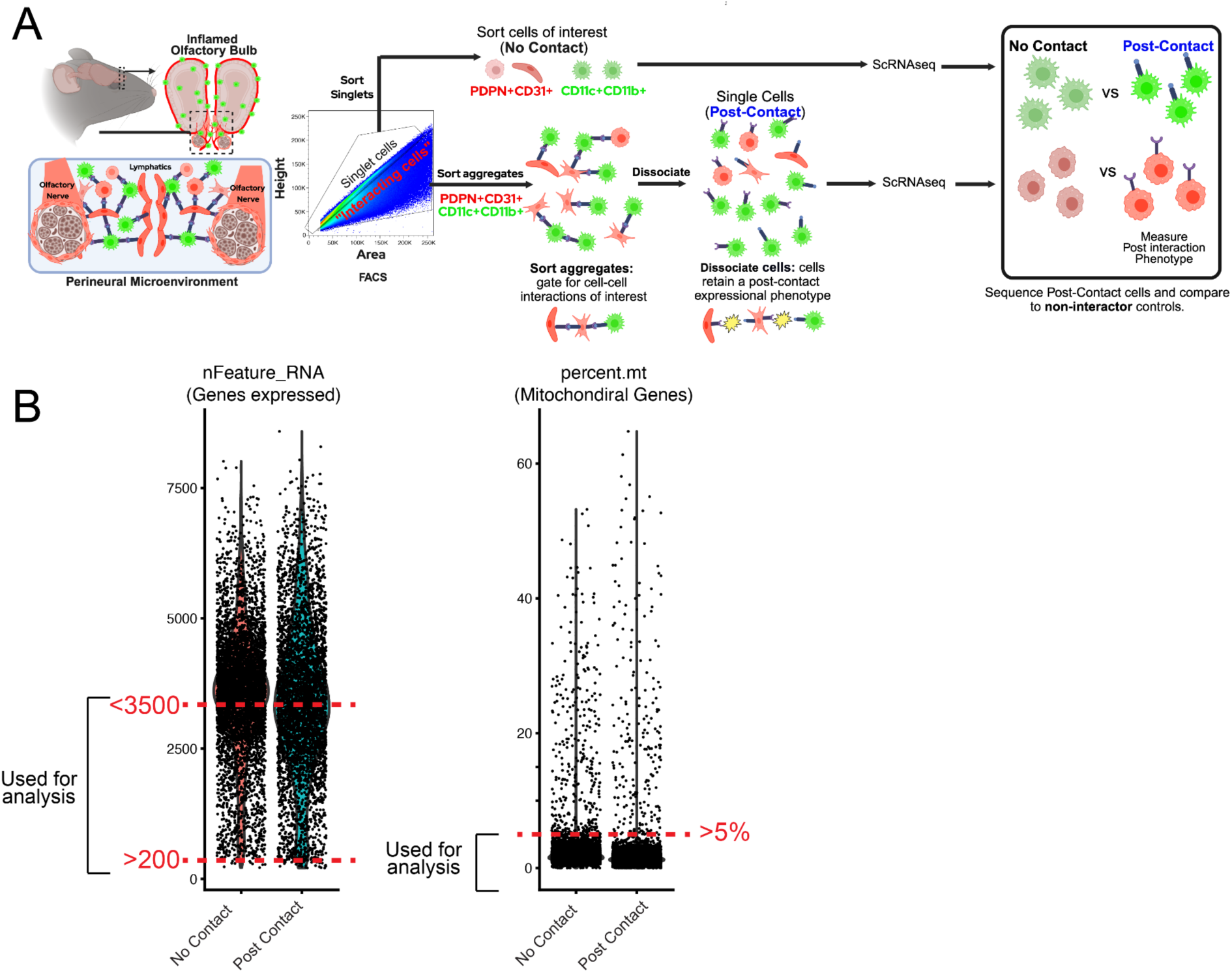
Strategy to isolate CD11b+CD11c+PDPN+CD11b+ “No contact” and “Post-Contact” populations. **(A):** Cartoon scheme outlining “PostContact-seq” sorting procedure of cribriform plate + Olfactory bulb tissue preparations from EAE 3.0. Sorted singlets from CD11c^+^CD11b^+^ (Myeloid cells) and CD31^+^PDPN^+^ (Meningeal-Lymphatic niche) gates were collected into a no-contact tube. Simultaneously, quadruple positive CD11c^+^CD11b^+^CD31^+^PDPN^+^ (Myeloid cell+Meningeal Lymphatic Niche) aggregating cells were sorted into the “interactor” tube. Prior to scRNAseq, both tubes underwent short liberase treatment to dissociate interacting cells and generate single cell suspensions of post-contact, and sequenced. **(B) :** Cutoffs for filtered cells for differential analysis using the following criteria: Cells have > 200 genes expressed, Cells have < 3,500 genes expressed, Cells have < 5% mitochondrial genes expressed. Strategy limits doublet aggregates at time of sequencing (High nFeatureRNA) and dead cells (High Percentage Mt).

**Supplemental Figure 2.**
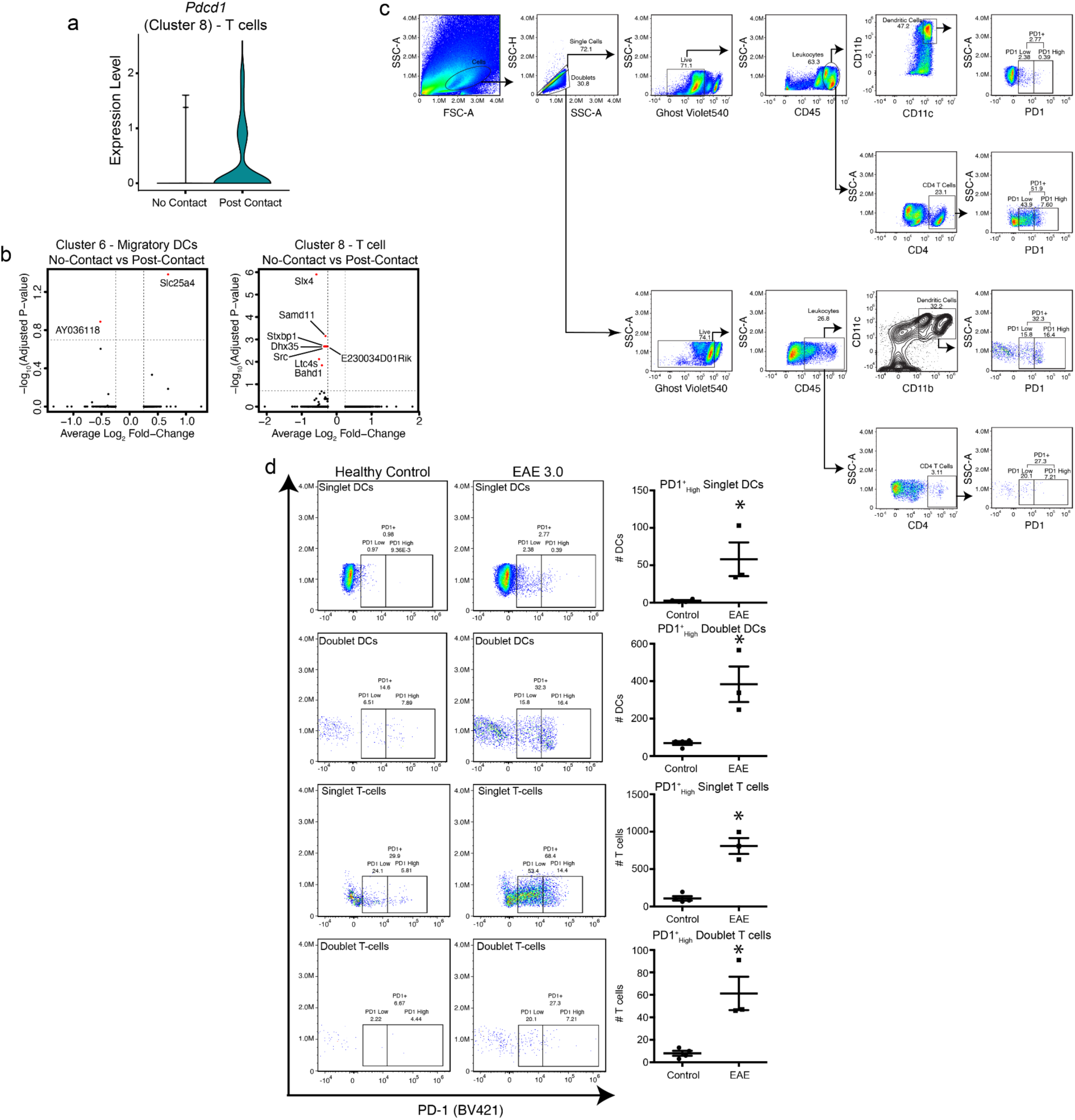
Analysis of PD-1^+^ immune subsets in isolated cribriform plate cell suspensions. **(A)** Violin plot showing *Pdcd1* (PD-1) expression in T cells (Cluster 8) stratified by inferred cell-cell interactions, with higher expression observed in post contact T cells. **(B)** Volcano plots of differential gene expression between post-contact and no-contact cells in Cluster 6 migratory dendritic cells (left) and Cluster 8 T cells (right) **(C)** Flow cytometry gating strategy for identifying singlet and doublet populations of CD45^+^ leukocytes, dendritic cells (CD11c^+^), and CD4^+^ T cells, followed by assessment of PD-1 expression across subsets. No liberase treatment given **(D)** Representative flow plots and quantification of PD-1^high^ singlet and doublet dendritic cells and T cells in healthy control and EAE day 3.0 mice. EAE mice show significantly elevated PD-1^high^ populations across all subsets, with the largest increase observed in doublet T cells, suggesting enhanced immune interaction and checkpoint activation during neuroinflammation. Singlet DCs (p=0.0322), Doublet DCs (p=0.0084), Singlet T cells (p=0.0005), Doublet T cells (p=0.0007). Unpaired two-tailed t test. Data are represented as mean ± SEM

**Supplemental Figure 3.**
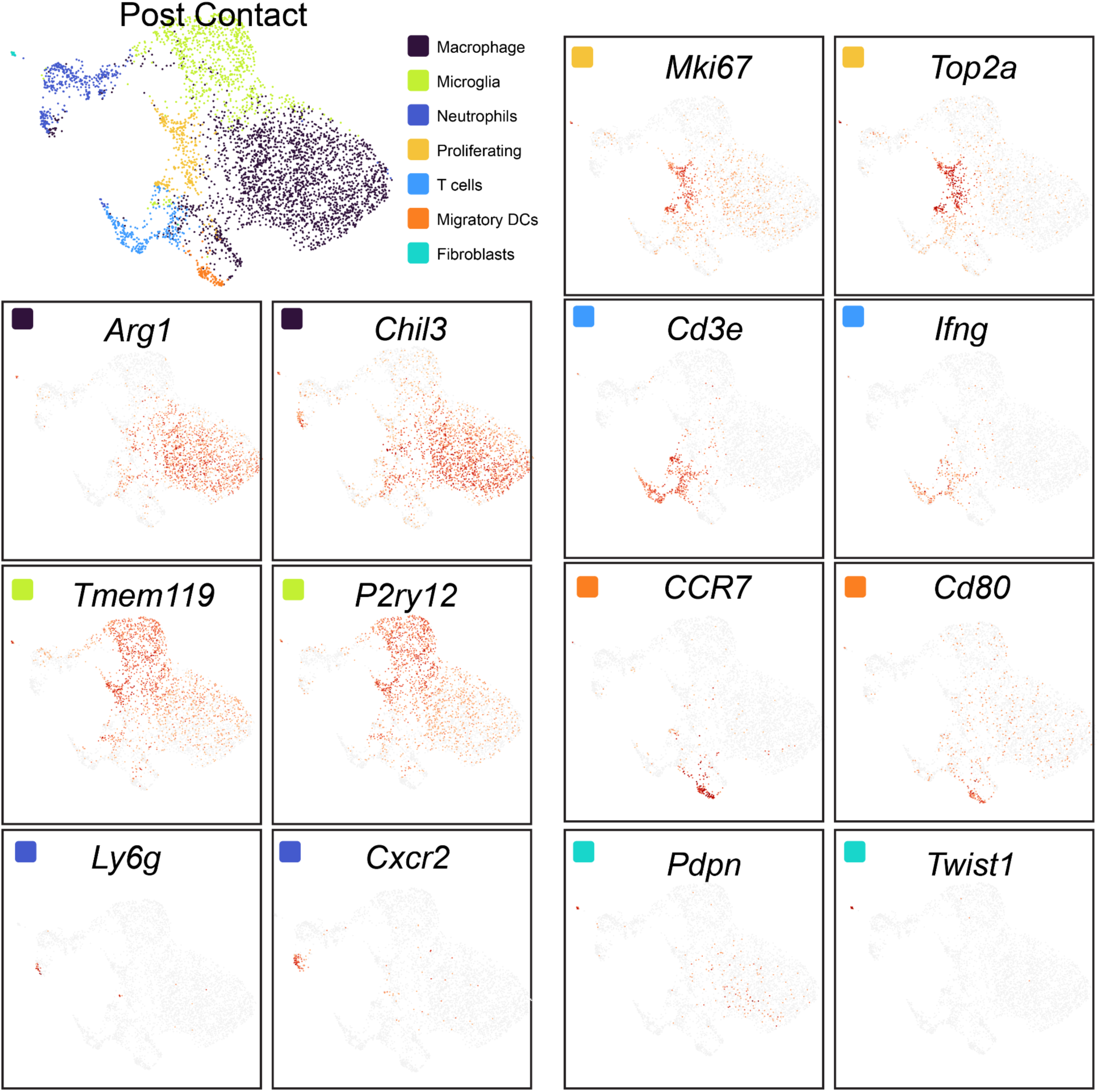
Cell cluster characterization of “post contact” CD11b^+^CD11c^+^ PDPN^+^CD31^+^ interacting cells. UMAP projection (top left) shows major post contact cell types isolated, including macrophages (purple), microglia (green), neutrophils (blue), T cells (orange), proliferating cells (light blue), migratory dendritic cells (yellow-orange), and fibroblasts (aqua). Feature plots highlight expression of *Arg1*, *Ly6c2*, *Ccr2*, *Chil3* (CHI3L3), and *Pdpn*, identifying infiltrating monocyte-derived, alternatively activated macrophages, distinct from T cells (*Cd3e*, *Ifng*), DCs (*CCR7*, *Cd80*), microglia (*Tmem119*, *P2ry12*), neutrophils (*Ly6g*, *Cxcr2*), and proliferating cells (*Mki67*, *Top2a*). Gene ontology enrichment analysis (bottom) demonstrates that these post contact macrophages are enriched for pathways in Fcγ receptor and complement receptor signaling, negative regulation of IL-12 production, homotypic cell-cell adhesion, and hemostasis.

**Supplemental Figure 4.**
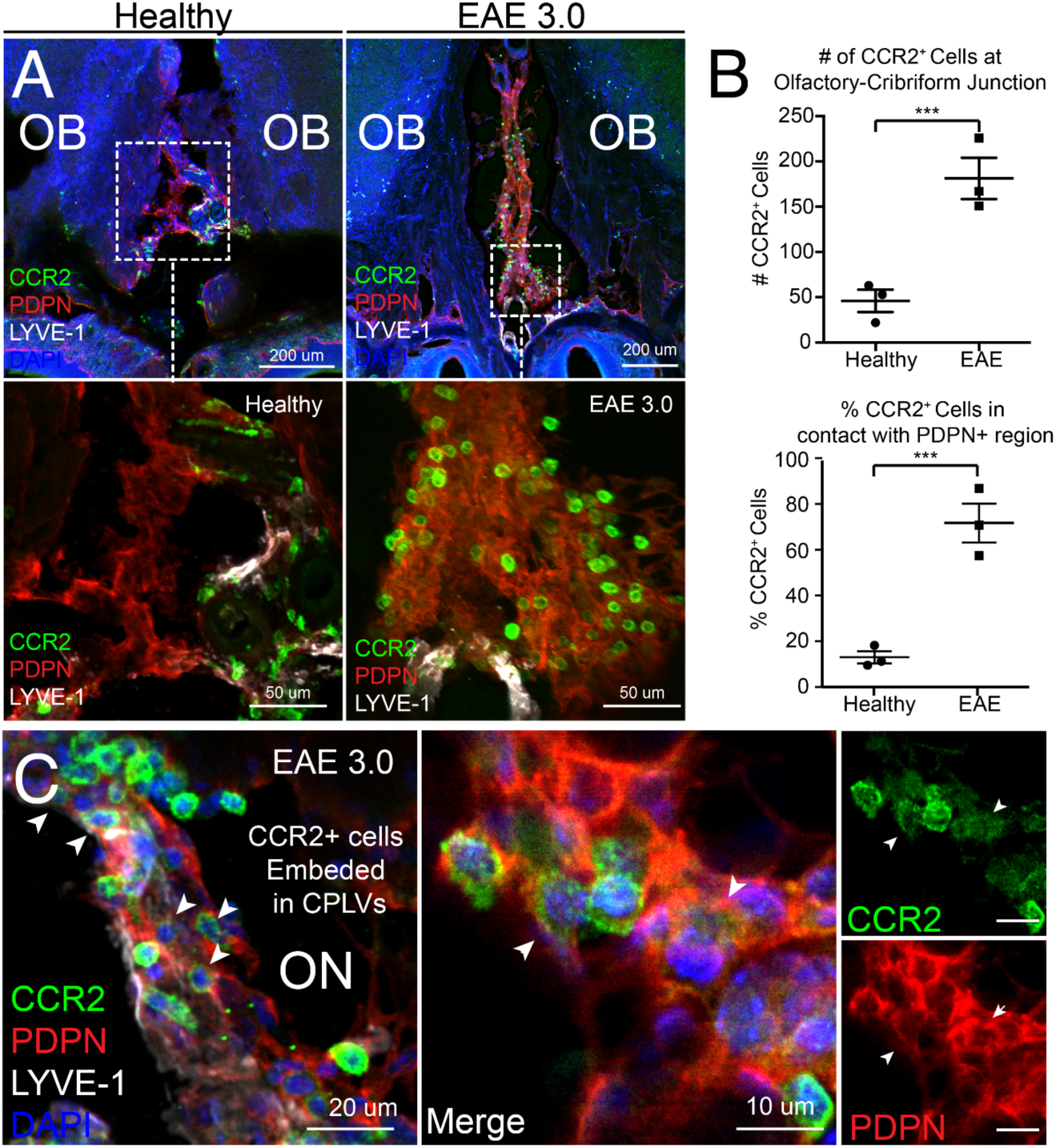
Accumulation of CCR2^+^ immune cells in PDPN^+^ cribriform regions during EAE. **(A)** Immunofluorescence images of healthy and EAE 3.0 olfactory bulbs (OBs), showing increased CCR2⁺ macrophage accumulation at the olfactory-cribriform junction. Insets show higher magnification of boxed areas with increased CCR2⁺ cell localization to PDPN-rich regions in EAE. **(B)** Quantification of CCR2⁺ cell numbers and their proportion in contact with PDPN-rich zones at the olfactory-cribriform interface. EAE mice show significantly more CCR2⁺ cells and higher PDPN-association. Unpaired two-tailed t test. Data are represented as mean ± SEM (p = 0.0027). **(C)** High-magnification image showing CCR2⁺ cells (green) embedded within cribriform plate lymphatic vessels (CPLVs, marked by LYVE-1, white) and adjacent to PDPN⁺ region (red) at the olfactory nerve (ON) interface in EAE 3.0 mice. Arrowheads indicate CCR2⁺ cells associated with CPLVs. Unpaired two-tailed t test. Data are represented as mean ± SEM (p = 0.0064)

**Supplemental Figure 5.**
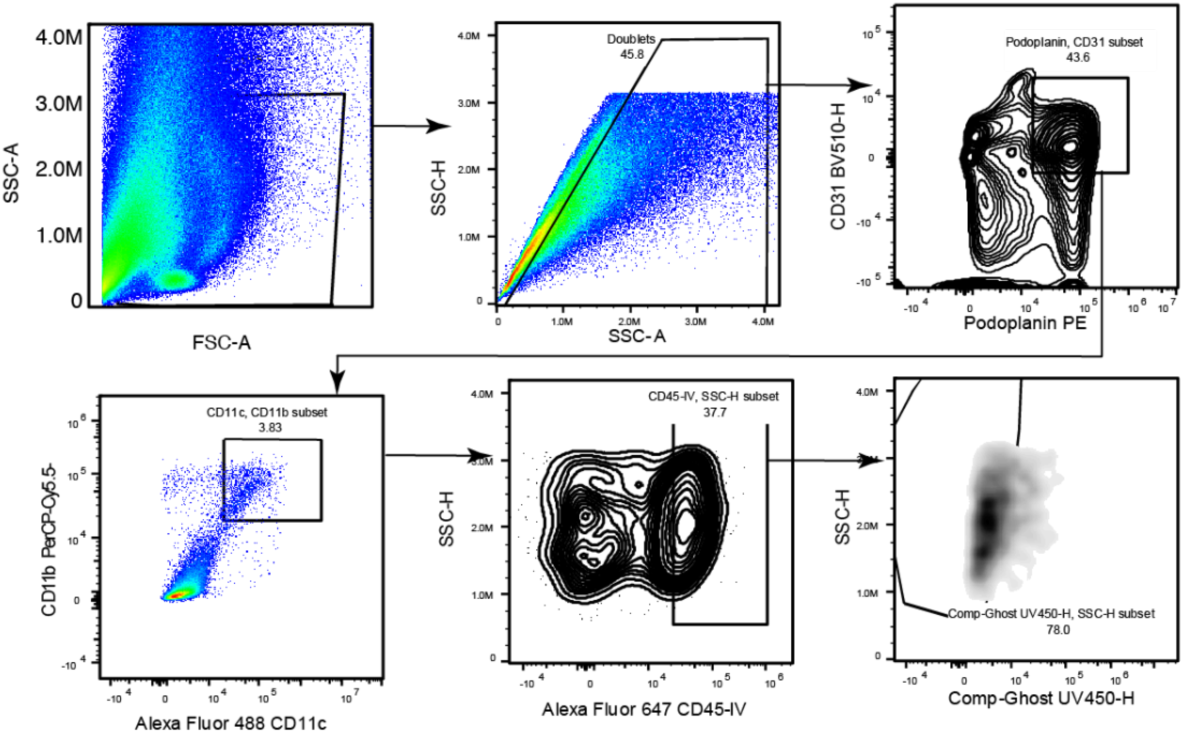
Gating strategy for CD45-IV+ interactors. Isolation of specific cell populations, beginning with size-based selection (FSC-A/SSC-A) and doublet inclusion. The hierarchy branches to identify endothelial and stromal subsets (CD31/Podoplanin) while simultaneously profiling myeloid compartments through specific surface markers (CD11c/CD11b). Finally, the analysis incorporates intravascular cell labeling (CD45-IV) and viability dye (Ghost UV450-H) to precisely quantify live leukocyte populations within the sample.

**Supplemental Figure 6.**
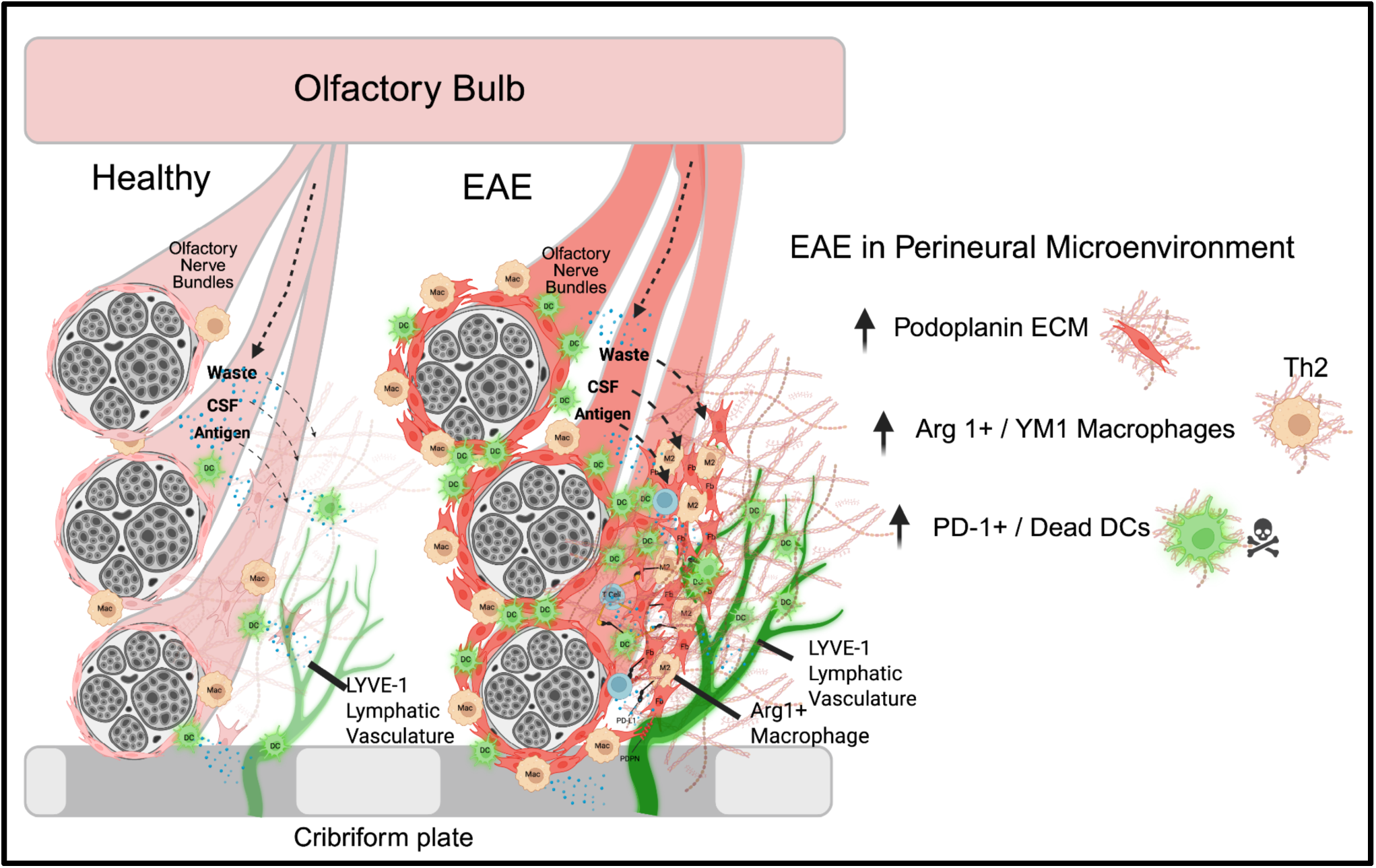
Graphical Summary. Perineural Microenvironment (PME) has Expanded Immune-Stromal-Lymphatic Niche During EAE. In the healthy state the PME of olfactory nerve bundles (ON) have lower levels of myeloid cells and lymphatics but still have access to draining cerebrospinal fluid (CSF), waste, antigen, and immune cells like DCs. During EAE neuroinflammation the PME becomes remodeled with higher myeloid cell accumulation within PDPN+ regions: fibroblasts, lymphatics, ECM, and macrophages. This creates an assembled immunoregulatory environment around olfactory nerve bundles where tolerogenic myeloid cells engage in cell-cell interactions alongside nerve bundles and cribriform lymphatics.

